# Fault-tolerant 3D reconstruction from 2D spatial proteomics sections

**DOI:** 10.64898/2026.06.23.733649

**Authors:** Zhaojun Zhang, Yuqi Tan, Michael Snyder, Garry P Nolan, Zongming Ma

**Affiliations:** Department of Statistics and Data Science, The Wharton School, University of Pennsylvania, PA, United States; Department of Microbiology and Immunology, Stanford University, CA, United States; Department of Pathology, Stanford University, CA, United States; Department of Genetics, Stanford University, CA, United States; Department of Statistics and Data Science, Yale University, CT, United States; Department of Biomedical Informatics and Data Science, Yale University, CT, United States

**Keywords:** 3D inference, Diffusion model, Generative AI, Multiplexed imaging, Spatial proteomics

## Abstract

Reconstructing 3D molecular volumes from sparsely sampled 2D tissue sections is limited by per-section marker dropout and tissue loss. We present 3D-Omics-Flow, a generative pipeline that jointly repairs damaged sections and interpolates between them at single-cell resolution. Across datasets spanning health and disease, 3D-Omics-Flow expands 3D spatial proteomics to practical sampling regimes, enabling atlas construction and downstream analysis from imperfect 2D section stacks.

---

Spatial proteomics measures protein abundance at single-cell resolution in intact tissue (1, 2), but tissue architecture extends in depth, and individual 2D sections capture only fragmented views of multicellular niches, vascular–immune interfaces, and invasive fronts. Dense serial sectioning or direct 3D spatial proteomics could recover this context, yet the cost and antibody-based nature of most platforms make routine 3D acquisition impractical (3). Reconstruction from sparsely sampled 2D sections is therefore an attractive alternative (4, 5), but in spatial proteomics, it is limited by two failure modes that are common in large-tissue workflows: section-specific biomarker dropout and tissue damage from tearing or folding. Existing methods fail to thoroughly address these practical challenges. SpatialZ (5) demonstrated sparse-3D reconstruction in spatial proteomics only at small scales and with undamaged 2D sections, and image-domain interpolation methods such as InterpolAI (6) are not designed for large *z*-axis gaps or high-plex marker panels. Here we present 3D-Omics-Flow, a generative pipeline that jointly repairs each observed 2D section by imputing dropped markers and restoring damaged tissue regions, and interpolates the restored stack at single-cell resolution, supporting downstream 3D analyses, including virtual sectioning, continuous-gradient profiling, 3D tissue microenvironment, and cell-cell interaction mapping (Fig. 1a).

**Figure 1.**
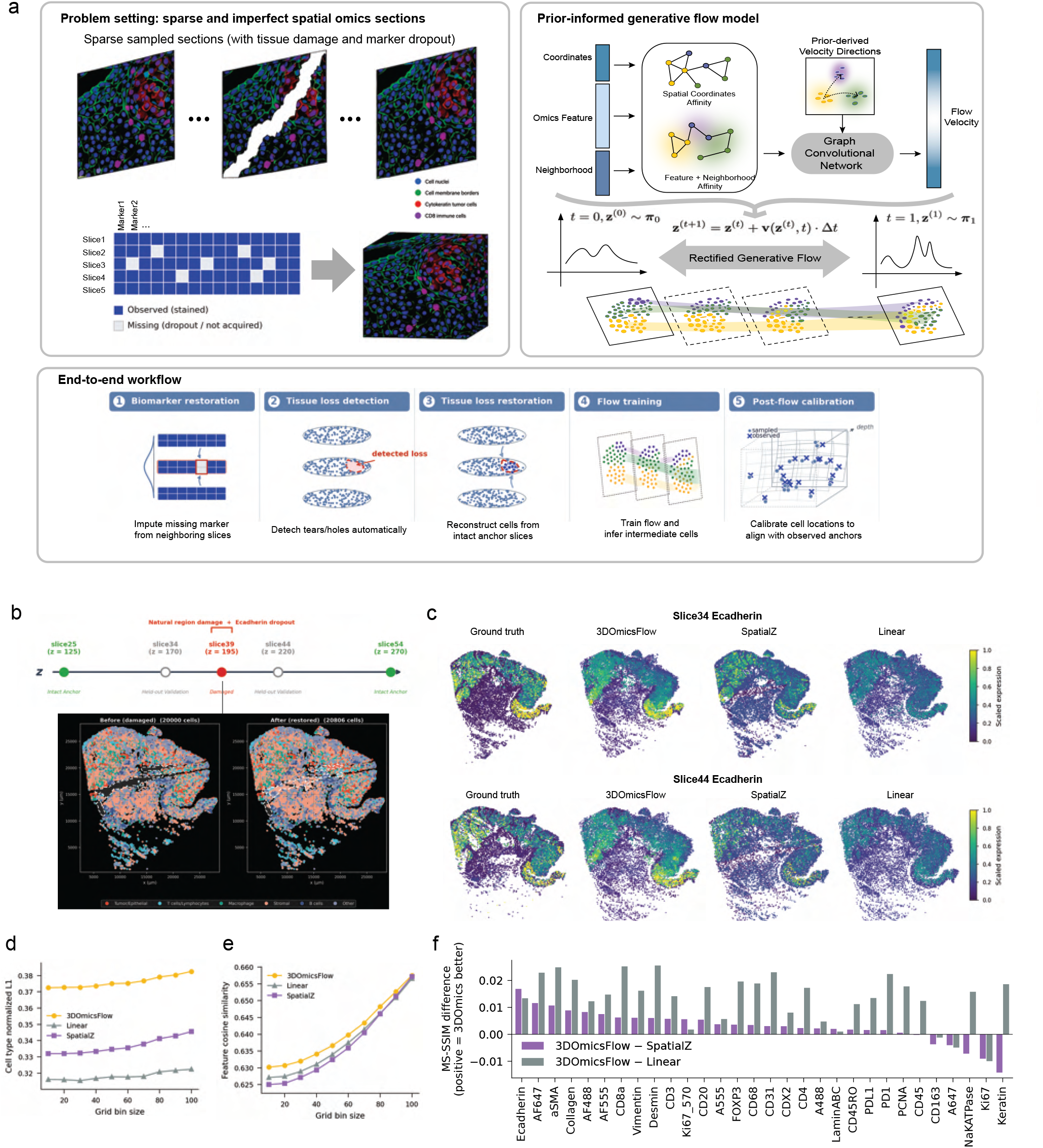
3D-Omics-Flow reconstructs a continuous 3D spatial-omics atlas from sparse, partially damaged 2D sections. **a**, Method overview. Top left: data format (sparse 2D sections with tissue damage, partial marker dropout, or both). Top right: prior-informed generative flow model. The model learns a velocity field over each cell’s spatial-molecular state and integrates it via rectified flow to infer intermediate cellular states between adjacent slices. Bottom: end-to-end workflow: (1) biomarker restoration; (2) tissue-loss detection; (3) tissue-loss restoration; (4) velocity-field encoding; (5) flow training; (6) post-flow calibration. **b**, Natural-damage benchmark on CRC proteomics. Three training inputs (slice25, *z* = 125 *µ*m and slice54, *z* = 270 *µ*m as intact anchors; slice39, *z* = 195 *µ*m carrying a damaged tissue region and E-cadherin dropout) and two held-out validation slices (slice34, *z* = 170 *µ*m; slice44, *z* = 220 *µ*m). Below: 2D scatter of slice39 before (left) and after (right) 3D-Omics-Flow restoration with the gap filled by inferred cells, colored by major cell type. **c**, Inspection of per-slice reconstruction at the two held-out validation sections, slice34 (top row) and slice44 (bottom row), for the damaged E-cadherin channel: ground truth, 3D-Omics-Flow, SpatialZ, and linear interpolation baseline (from left to right). **d**,**e**, Spot-level metrics across bins on the held-out validation slices as a function of grid bin size 10–100 *µ*m; higher is better for both metrics. **d**, cell-type-normalized *L*_1_ score; **e**, feature cosine similarity score. **f**, Per-marker MS-SSIM difference on the validation slices: Improvements by 3D-Omics-Flow over SpatialZ (purple) and over Linear (gray). Positive values indicate 3D-Omics-Flow reconstructs the marker more faithfully than the respective benchmarking method.

We benchmarked 3D-Omics-Flow on a publicly available colorectal cancer (CRC) spatial proteomics cohort acquired by serial-section CyCIF (7) to simulate an experimental scenario exhibiting both structural damage and missing staining: out of the three training slices (#25, #39, and #54), one (#39) had a *natural* tissue damage (Fig. 1b, lower-left, dashed region) and a *simulated* E-cadherin marker dropout. After training 3D-Omics-Flow and 3D reconstruction based on the three training slices, we evaluated the predictions on two held-out validation slices (#34, #44) (Fig. 1b) which were shielded from the entire model fitting and inference procedure, including but not limited to preprocessing, normalization, feature engineering, clustering, label assignment, and parameter tuning. See Materials & Methods for details. 3DOmics-Flow restored the tissue loss on slice #39 (Fig. 1b, lower-right), and reconstructed a 3D volume.

Qualitatively, when the reconstructed volume was sectioned at the *z*-coordinates of the two validation slices, the 3DOmics-Flow output reproduced the ground-truth spatial organizations of E-cadherin, whereas SpatialZ (5) and naïve linear interpolation (Linear) failed to recover these patterns with comparable fidelity (Fig. 1c). See, for instance, the over-prediction around the middle-left region in both validation slices by SpatialZ. Quantitatively, 3D-Omics-Flow outperformed SpatialZ and Linear by a sizable margin in cell type annotation fidelity across spatial resolution levels: cell-type normalized-*L*_1_ similarity scores (Fig. 1d) based on 3D-Omics-Flow predictions were consistently higher (with ~12–19% relative improvements) across spot sizes from 10 to 100 *µ*m. Moreover, 3D-Omics-Flow consistently outperformed SpatialZ and Linear in feature prediction accuracy across different spatial resolutions in both overall (Fig. 1e, measured in cosine similarity between spot-level predicted and true feature vectors across spot sizes between 10 and 100 *µ*m) and per-feature (Fig. 1f, better than both in 25 out of the 30 measured biomarkers, measured in per-feature MultiScale Structural Similarity, i.e., MS-SSIM, scores) sense. Notably, the largest per-feature improvement over SpatialZ occurred exactly on E-cadherin (Fig. 1f), the dropout marker on the middle training slice. Finally, the computation time cost for 3D-Omics-Flow is significantly smaller and scales much better than that of SpatialZ (Supplementary Fig. S1). These improvements resulted from a carefully-designed workflow and computational strategy for dropout marker imputation and tissue loss restoration tailored to the present 3D volume prediction setting, which is not addressed by any state-of-the-art sparse-3D reconstruction method.

Across spatial proteomics (CyCIF (7), INSIHGT (8)) and spatial transcriptomics (OpenST (9)), 3D-Omics-Flow predictions matched ground-truth spatial structure on held-out validation slices (Supplementary Figs. S2 and S3), retained community-level geometry and depth-resolved abundance dynamics on the ground-truth volumetric mouse hypothalamus dataset (Supplementary Fig. S4), and preserved molecular fidelity at the level of pairwise biomarker–biomarker correlations and 3D ligand–receptor scores (Supplementary Fig. S5). The tissue loss restoration capacity of 3D-Omics-Flow on 2D spatial transcriptomics stacks was demonstrated on a lymph node OpenST dataset in Supplementary Fig. S3 and S6.

Building on these proof-of-concept results, we next applied 3D-Omics-Flow to a larger CRC stack (also from (7), but non-overlapping with that used in Fig. 1) with more extensive staining corruption and per-section damage generated by simulation. The stack comprised 11 training slices spanning ~460 *µ*m along the *z*-axis, with three section types: intact, combined damage (tissue loss + feature dropout), and feature-dropout-only damage (Fig. 2a, green/red/orange sections, respectively). We simulated rectangular tissue loss regions to mimic the realistic scenario in which a researcher is aware of tissue loss within such a bounding box while pinning down the exact loss boundary is difficult. 3D-Omics-Flow assembled a continuous 3D atlas from these training slices alone. When evaluated on held-out validation sections (Fig. 2a, uncolored sections), 3D-Omics-Flow predicted molecular structures with high fidelity (Fig. 2b). It also uncovered the 3D cellular landscape (Fig. 2c), restoring cross-slice continuity of cell populations that appear fragmented when viewed in isolated 2D sections.

**Figure 2.**
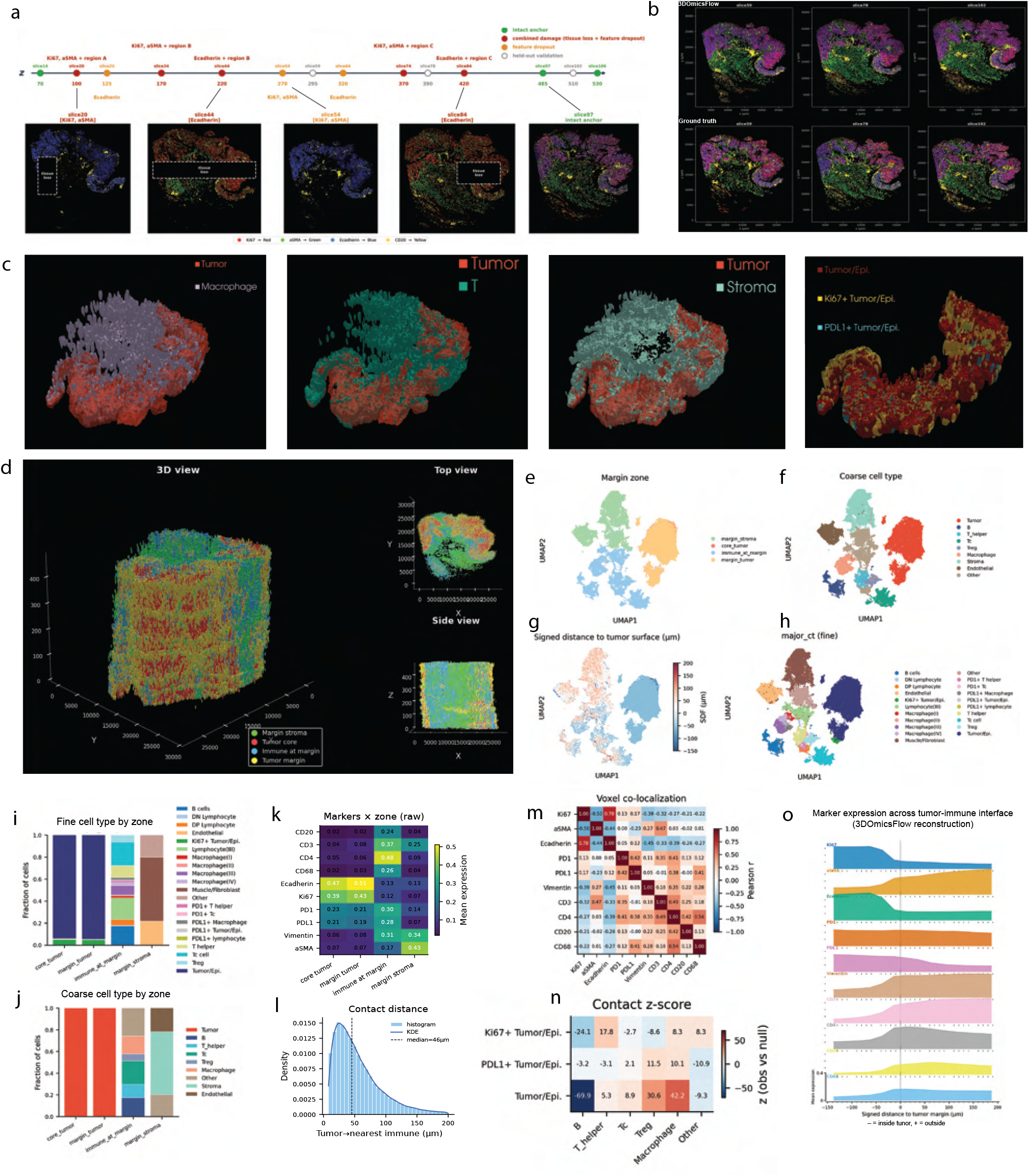
3D-reconstructed CRC atlas reveals continuous tumor-immune architecture inaccessible from sparse 2D sections. **a**, Combined-damage design on the CRC proteomics cohort. 11 training slices spanning *~*460 *µ*m along the *z*-axis, with three section types: intact, combined damage (tissue loss + feature dropout), and feature-dropout-only damage, corresponding to green, red, and orange sections. Three uncolored sections were held-out validation sections and were shielded from the entire training and 3D reconstruction process. Below: examples of damaged input slices. **b**, Comparison of reconstructed 3D volume with ground truth at *z*-coordinates of held-out validation slices: side-by-side feature overlays, 3D-Omics-Flow vs. ground truth. **c**, Components of the 3D volume reconstructed by 3D-Omics-Flow colored by cell-group combinations: Tumor+Macrophage, Tumor+T, Tumor+Stroma, and Ki67^+^ vs. PD-L1^+^ tumor/epithelial subtypes. **d**, Tumor-immune invasive-margin zones (tumor core, tumor margin, immune at margin, margin stroma) defined by a signed-distance field over a dilated tumor voxel mask (25 *µ*m 3D grid). Left: 3D oblique view; Right: top (XY) and side (XZ) projections. **e-h**, UMAP visualizations of observed and virtual cells in 3D-Omics-Flow-reconstructed volume colored by invasive-margin zone (**e**), coarse cell group (**f**), signed distance to the tumor surface in *µ*m (**g**) and fine cell subtype (**h**). **i**,**j**, Zone composition: stacked-bar fractions per zone for fine cell subtype (**i**) and coarse cell group (**j**). **k**, Mean expression of canonical markers (rows) across invasive-margin zones (columns). See Supplementary Fig. S7 for further details, including differentially-expressed markers in core tumor vs. margin tumor zones. **l**, Distribution of nearest tumor → immune Euclidean distances for margin-tumor cells. **m**, Voxel-level (25 *µ*m 3D grid) Pearson correlation between markers across the full 3D reconstruction. **n**, Tumor-immune contact *z*-scores within 60 *µ*m of tumor boundary (tumor/epithelial cell subtypes vs. immune cell types) against a label-shuffled null (*n* = 200 permutations). **o**, Marker expression as a function of signed distance to the tumor surface (negative = inside tumor, positive = outside tumor).

Within the reconstructed volume, a signed-distance field from each cell to the tumor surface delineated four invasivemargin zones (tumor core, tumor margin, immune at margin, and margin stroma; Fig. 2d) and revealed a continuous phenotypic gradient from tumor core to margin stroma (Fig. 2e-h). Cell-type composition differed across zones, with peak diversity in the immune at margin and margin stroma (Fig. 2i,j). CD3^+^/CD4^+^ T-cells were largely confined to the margin and excluded from the tumor core, coincident with an *α*-SMA-high stroma at the interface, defining an immune-excluded phenotype (Fig. 2k). Marker gradients along the signed distance to the tumor surface revealed PD-L1 expression rising sharply at the interface and a ~100-*µ*m-wide CD3^+^/CD4^+^ T-cell belt accumulating at the stromal boundary (Fig. 2o), consistent with combined stromal sequestration and PD-L1-associated exclusion of T-cells from the tumor core. We quantified tumor– immune spatial relationships in 3D via tumor-to-immune nearest-neighbor distances and voxel-level marker correlations (Fig. 2l,m). Neighborhood enrichment within 60 *µ*m of interface tumor cells revealed subtype-specific immune niches: Ki67^+^ tumor cells were enriched for T-helper and macrophage contacts, PD-L1^+^ tumor cells for Treg and macrophage contacts, and bulk tumor/epithelial subtypes were depleted of B-cell contacts (Fig. 2n), indicating that proliferative and immune-evasive tumor states occupy distinct local microenvironments. Transferring these 3D-derived zone labels onto held-out validation slices uncovered statistically significant margin tumor vs. core tumor differences in proliferation and immune markers that the corresponding sparse 2D zonation failed to detect (Supplementary Fig. S7). Notably, the paradigm of 3D-reconstructionby-training-slices plus *p*-value-calculation-on-held-out-2Dvalidation-slices adopted here provides a generic way for testing differentially-expressed biomarkers in different 3D neighborhoods with statistical rigor, which can be further generalized to other downstream inference tasks.

In sum, 3D-Omics-Flow addresses a core practical limitation of 3D spatial-omics reconstruction: the combination of sparse *z*-axis sampling, biomarker dropout, and physical tissue damage that routinely arises in large-scale serial-section workflows. Unlike deep-learning approaches (e.g., (10, 11)) that depend on large modality-specific pretraining datasets, 3D-Omics-Flow is designed for data-scarce settings, making it well-suited to studies of rare tissues and to modalities for which dense 3D atlases remain prohibitively expensive. On a CRC CyCIF cohort, 3D-Omics-Flow recovered held-out 3D structure under combined biomarker dropout and tissue damage, whereas SpatialZ (5) did not. 3D-Omics-Flow generalized to clean dense 3D proteomics (INSIHGT mouse hypothalamus) and to spatial transcriptomics with (simulated) tissue damage (OpenST metastatic lymph node). The reconstructed CRC volume enabled 3D tumor-immune analyses inaccessible from standalone 2D planes: continuous molecular gradients across the invasive front, voxel-level marker correlations, and tumor-immune contact enrichment within a fixed spatial radius. Although 3D-Omics-Flow is designed for spatial proteomics, future iterations could more explicitly leverage multi-modal integration, broadening 3D spatialomics analysis from a few well-resourced platforms to atlas-scale studies across diverse spatial modalities.

## Materials & Methods

### Preliminaries for 3D-Omics-Flow

#### Notation

Let *k* ∈ { 1,…, *K*} index slices, with *n*_*k*_ denoting the total number of cells on slice *k* and *i* ∈ { 1,…, *n*_*k*_} indexing cells within slice *k*. The raw omics feature matrix for slice *k* is 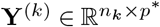 (where *p*^∗^ is the number of distinct features measured across all slices), and the total number of cells is 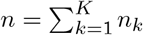. When marker dropout is present in slice *k*, each corresponding column in **Y**^(*k*)^ is filled with zeros. After preprocessing and dropout feature restoration, the representation feature matrix for slice *k* is denoted by 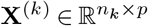, where *p* is the representation feature dimension. The raw cell locations in slice *k* are stored in the coordinate matrix **S**^(*k*),raw^. After spatial alignment of slices detailed below, the post-alignment cell coordinates on slice *k* are denoted by **S**^(*k*)^. In what follows, we let 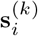 and 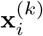 be the (postalignment) coordinate vector and representation feature vector for cell *i* in slice *k*, respectively.

#### Spatial Alignment of Slices

The raw coordinates of serial 2D sections are not directly comparable across slices. Thus, we co-register adjacent slice pairs through a rigid initialization followed by a non-rigid refinement. Given a pair of consecutive slices (e.g., slices *k* and *k* + 1), we first rasterize each slice’s cell coordinates to a 2D density image and recover a rigid initialization between target slice *k* and source slice *k* + 1 via a two-pass rotation grid search (coarse 10^°^ step over [ −180^°^, 180^°^), followed by a refined 1^°^ step in a ±10^°^ window around the coarse optimum). At each candidate rotation *θ* (applied to the source density), we select the translation (Δ*a*, Δ*b*) that maximizes the normalized cross-correlation between the target density *H*^(*k*)^ and the rotated source density 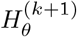,

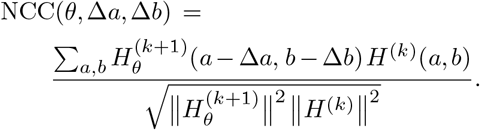

Using rasterized density images (rather than centroid matching) keeps the rigid initialization robust to missing tissue regions, since absent cells contribute zero to the density images and therefore do not bias the correlation peak. In the second stage, we pass this rigid initialization to STalign (12), which further refines the registration with a non-rigid deformation. The resulting deformation is applied to every cell of slice *k* + 1, and the aligned coordinates **S**^(*k*+1)^ replace **S**^(*k*+1),raw^ for all subsequent preprocessing and modeling. This procedure is applied iteratively for *k* = 1,…, *K* − 1, with slice 1 serving as the anchor whose raw coordinates define the canonical frame: **S**^(1)^ ≡ **S**^(1),raw^. At each iteration, slice *k* + 1 is registered to the already-aligned **S**^(*k*)^, and the resulting **S**^(*k*+1)^ then becomes the target for registering slice *k* + 2 at the next iteration.

### Pipeline Overview

3D-Omics-Flow reconstructs a continuous 3D omics volume from a stack of *K* 2D slices through five sequential steps that follow a *“restore-first, generate-later”* principle. The pipeline assumes that the top and bottom slices of the stack do not have tissue damage and that at any specific point in the shared coordinate system, only a minority of slices suffer tissue damage. Steps 3 and 4 share a common generator, the two-slice building block Eq. (1) (formalized in Interlude below), which produces a continuous interpolation between two complete slices that bracket an unseen interval. The first three steps restore each input slice to a complete structurally contiguous state: Step 1 (biomarker restoration and preprocessing) imputes all missing molecular features across slices by taking into account both spatial and feature-level dependence, then normalizes features, computes a CAST graph embedding, standardizes coordinates, and curates spatial-community and cell-type priors for downstream modeling; Step 2 (tissue loss detection) identifies, without manual annotation, which slices contain missing spatial regions, locates each such region, and assigns each damaged slice an intact anchor pair (the nearest slices without tissue loss above and below); Step 3 (tissue loss restoration) evaluates the building block between the assigned anchor pair to reconstruct cells in each detected missing region. The remaining two steps perform cross-slice 3D interpolation and post-flow calibration: Step 4 (pairwise generation and cross-interval assembly) applies the building block to each adjacent slice pair of the restored stack to obtain matched-region trajectories and unmatched-region samples, and stitches the per-interval predictions at the observed slice levels; and Step 5 (post-flow calibration) aligns the generated coordinates to any additional positional information. When the stack has *K >* 2 slices, Step 1 is applied per slice; Step 2 pools the stack to compute a cross-slice reference density before flagging per-slice damage masks; Step 3 is applied per damaged slice using its anchor pair from Step 2; Step 4 instantiates the building block pairwise on each adjacent interval of the restored stack; and Step 5 calibrates the assembled stack globally in one shot.

### The 3D-Omics-Flow Algorithm

#### Step 1: Biomarker Restoration

For slice *k*, let ℱ_*k*_ be the column index set of measured biomarkers in **Y** ^*k*)^, and 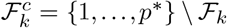 collect the indices of the dropout features. Define the set of shared features 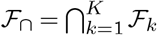, which is observed across all slices and serves as the bridge for cross-slice information transfer. In addition, we require that 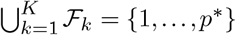. The goal of Step 1 is to impute the missing columns of each **Y**^(*k*)^ by leveraging both the shared features and the spatial tissue context.

##### Within-slice Spatial Graph Construction

Let *κ* be a positive integer (default *κ* = 15). For each slice *k*, we construct a *κ*-nearest-neighbor graph on the spatial coordinates **S**^(*k*)^. For an edge connecting cells *i* and *j*, the edge weight is

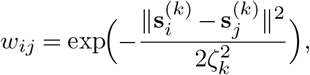

where *ζ*_*k*_ is set to the median *κ*-nearest-neighbor distance within slice *k*. The edge weight matrix is then symmetrized by assigning the Gaussian weight to both directed edges whenever *j* is a *κ*-NN of *i* (so *W*_*ij*_ = *W*_*ji*_ if *j* ∈ *κ*NN(*i*) or *i* ∈ *κ*NN(*j*)) and row-sum-normalized to yield a row-stochastic adjacency matrix **A**^(*k*)^. The slice-specific adjacency matrices are assembled into a block-diagonal matrix **A** = blkdiag(**A**^(1)^, …, **A**^(*K*)^) ∈ R^*n×n*^, which preserves slice isolation.

##### Spatial Context Augmentation

For each slice *k*, let 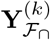 be the *n*_*k*_-by|ℱ_∩_| matrix that only retains the columns corresponding to the biomarkers measured by all slices. Let 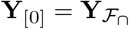 be the *n*-by-|ℱ_∩_| matrix obtained from stacking 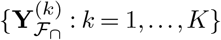 in order. For *h* = 1, …,*H* (default *H* = 2), define the *h*-hop weighted neighborhood mean

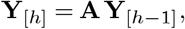

which averages expressions over progressively larger spatial neighborhoods. In addition, define three spatial statistics.

1. *Neighborhood dispersion*. For each cell *i* and feature *f*,

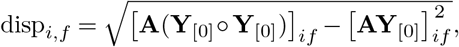

where ° denotes element-wise multiplication. This statistic captures local expression heterogeneity.
2. *Local cell density*. Within each slice *k*, for each cell *i* we compute the inverse mean distance to its *κ*_dens_ (default is 15) nearest spatial neighbors:

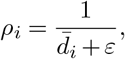

where 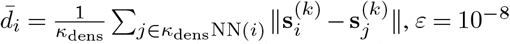
3. *Spatial gradient*. We define directional derivatives along the canonical planar axes as ∂**Y**_[0]_*/*∂*s*_1_ and ∂**Y**_[0]_*/*∂*s*_2_, which are estimated via weighted graph differences and capture spatial trends in expression.

We define the augmented descriptor of each cell by concatenating the preceding statistics

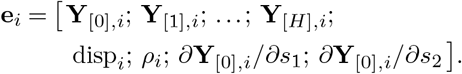

##### Z-Distance-Weighted Ridge Transfer

Consider a recipient slice *k*_rcv_ with missing-feature set 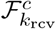; cells in all other slices serve as sources, weighted by *z*-proximity to *k*_rcv_:

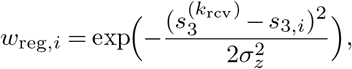

where *σ*_*z*_ is the median inter-slice distance. We then fit a weighted ridge regression of the source-cell target features on the augmented descriptor:

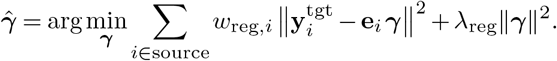

where 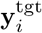 is the target-feature vector of source cell *i* for the recipient slice’s missing features, and *λ*_reg_ is chosen by cross-validation on held-out shared features by maximizing mean per-feature Pearson correlation. Predicted values populate the missing columns of 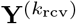, leaving observed entries unchanged.

##### Preprocessing

The biomarker-restoration step above produces a complete 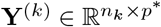 with all dropout columns populated. We complete Step 1 by executing the following preprocessing sub-steps in order.

1. *Modality-specific feature normalization*. For transcriptomics data (e.g., OpenST Human Lymph Node dataset), we apply library-size normalization, a log1p (*x* → log(1 + *x*)) transform, and per-feature scaling computed after pooling cells across slices. For proteomics data (e.g., CyCIF Human Colorectal Cancer and INSIHGT Mouse Hypothalamus datasets), we clip each feature to its pooled [0.005, 0.995] quantiles, linearly rescale to [0, 1], and then apply per-feature *z*-normalization.
2. *Representation feature matrix*. The processed feature matrices from all slices are passed to CAST (13) to obtain a graph embedding. For each slice *k*, the resulting embedding 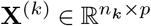, the representation feature matrix introduced in the Notation paragraph, serves as the feature matrix in the rest of the 3D-Omics-Flow pipeline unless stated otherwise.
3. *Coordinate normalization*. The coordinate matrix **S**^(*k*)^ is z-scored before downstream modeling, and generated coordinates are transformed back to the original scale.
4. *Spatial-community-cell-type priors*. We use cell-type and spatial-community annotations as prior information to guide flow model training. When such annotations are not provided by human experts, we apply Banksy (14) with *λ*_Banksy_ = 0.8 across all slices, using both spatial coordinates and omics features, to identify *m*_*q*_ spatial communities, and we apply Leiden clustering across all slices at the default resolution (1.0), based on similarity in omics features, to obtain *m*_*c*_ celltype clusters.

#### Step 2: Tissue Loss Detection

Whereas Step 1 imputes missing entries in **Y**^(*k*)^, Step 2 operates entirely on the post-alignment cell coordinates **S**^(*k*)^ to produce three outputs that drive Step 3: a damage mask ℳ_*k*_ ⊂ ℝ^2^ for each affected slice, a partition of the stack into damaged and healthy slices, and an anchor-pair map *k* ↦ (*k*_−_, *k*_+_) for each damaged slice. The detection exploits the volumetric continuity of tissue across serial sections: a location that is empty on one slice but occupied on most of the others is flagged as missing.

##### Per-Slice Rasterization

We overlay all *K* slices on a common 2D grid in the post-alignment (*s*_1_, *s*_2_) plane, indexed by integer coordinates (*a, b*). The grid spans the union bounding box of all *K* post-aligned slices, tiled with square bins of edge length 1*/*100 of the longer-axis range. For each slice *k* we compute the cell count *ν*^(*k*)^(*a, b*) and a Gaussian-smoothed per-pixel density *D*^(*k*)^(*a, b*) (Gaussian bandwidth *σ*_KDE_ = 1.0 pixels, applied via `scipy.ndimage.gaussian_filter`).

##### Cross-Slice Reference

For each grid cell (*a, b*) we define a reference density that summarizes what tissue should be there if a slice were intact:

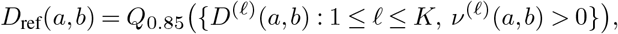

where *Q*_*p*_(·) denotes the *p*-quantile of a finite collection of real values; equivalently, *D*_ref_(*a, b*) is the 85th percentile of *D*^(*ℓ*)^(*a, b*) across slices with tissue at (*a, b*). We restrict attention to grid cells where *ν*^(*ℓ*)^(*a, b*) *>* 0 for at least two slices, which is reasonable under our assumption of complete tissues at the top and bottom of the stack.

##### Damage Mask Construction

A grid cell on slice *k* is a candidate for missing tissue if it contains *no cells at all* (*ν*^(*k*)^(*a, b*) = 0) yet the reference density there exceeds a threshold (*D*_ref_(*a, b*) *> τ*_*ρ*_) (default *τ*_*ρ*_ = 0.0375 · max_(*a′,b′)*_ *D*_ref_(*a*′, *b*′)); requiring strict emptiness, rather than just low density, ensures we flag true holes and not merely sparse regions. Candidate pixels are grouped into connected components and cleaned with a light morphology pipeline: binary closing with disk radius 3 pixels (to bridge single-pixel gaps), no binary opening, and no aggressive erosion. Components smaller than 5 pixels or smaller than 0.1% of the tissue envelope area are discarded, and adjacent components whose centroids lie within 1% range of the tissue diameter are merged, yielding the damage mask ℳ_*k*_ ⊂ ℝ^2^. A slice is labeled *damaged* when its total flagged area exceeds 1% of its footprint; the remaining slices are labeled *healthy* and serve as anchors for restoration in Step 3. For each damaged slice we record the nearest healthy slices above and below along *s*_3_ as its anchor pair; consecutive damaged slices that share the same pair are grouped together. Step 2 thus passes {ℳ_*k*_}, the damaged/healthy partition, and the anchor-pair map *k* ↦ (*k*_−_, *k*_+_) to Step 3.

##### Interlude: The Two-Slice Building Block

Before describing Step 3, we introduce the two-slice building block, a generator that produces a continuous interpolation between two complete slices that bracket an unseen interval. The building block is invoked in two contexts in 3D-Omics-Flow: (i) by Step 3 (described next), applied between two anchor slices to populate the missing region of a damaged slice at its depth between the anchors; and (ii) by Step 4 (described later), instantiated pairwise across all adjacent slice pairs of the restored stack to generate the 3D volume.

Suppose we are given two complete slices *D*_1_ and *D*_2_ that bracket an unseen interval, with no missing channels and no missing regions. We use a depth parameter *t* ∈ [0, 1] to index the relative position between the two slices (*t* = 0 corresponds to slice 1 and *t* = 1 to slice 2).

We rely on the two observed omics slices to obtain two different types of prior information on the unseen 3D volume between them: spatial communities and cell-type prototypes. As described in Step 1’s Preprocessing, community annotations are obtained by applying Banksy (14). Cell-type prototypes are obtained as the per-cluster average expression of Leiden clusters of the omics features.

We then train a community-specific, cluster-guided flow model that learns joint state trajectories (spatial coordinates, cell features, and neighborhood-feature context) between the matched regions of two slices. The velocity of the flow model is parameterized by a graph convolutional network (GCN). Together, these components enable controlled generation of cellular state trajectories conditioned on the prior information while preserving smooth interpolation between matched regions.

Concretely, the building block defines a procedure

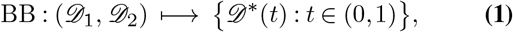

where, for *k* ∈ {1, 2, *D*_*k*_} comprises the post-preprocessing feature matrix **X**^(*k*)^, the post-alignment coordinates **S**^(*k*)^, and the spatial-community and cell-type annotations from preprocessing; each output virtual slice *D*^∗^(*t*) comprises a feature matrix **X**^∗^(*t*) and coordinates **S**^∗^(*t*), formed as the union of the matched-region trajectories generated by the flow model and the unmatched-region samples generated by the bin-level birth-death process.

Implementation details (initial matching and partitioning, cell state encoding, flow training, generative sampling, and the bin-level birth-death process for unmatched regions) are deferred to the later subsection titled “The Two-Slice Building Block: Components and Training”.

#### Step 3: Tissue Loss Restoration

For each damaged slice *k* with damage mask ℳ_*k*_ and anchor pair (*k*_−_, *k*_+_) from Step 2 (where 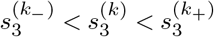, not necessarily immediate neighbors), we restore the cells inside ℳ_*k*_ by evaluating the building block Eq. (1) between the two anchors. The building block here is trained on the anchor pair 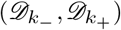, separately from the adjacent-interval instances used in Step 4.

##### Flow-Based Virtual Cell Generation

Concretely, we evaluate 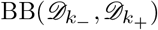 at the proportional depth

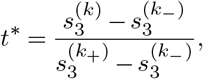

and set 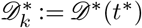; cells of 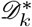 inside ℳ_*k*_ are the restoration candidates. The restored slice *k* then comprises the observed cells outside ℳ_*k*_ together with these restoration candidates. The candidate *z*-coordinates are rescaled to slice *k*’s level 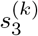 via the interval-specific rescale formula of Step 4 applied to the anchor interval (*k*_−_, *k*_+_).

By resolving all damaged slices, i.e., first restoring missing features (Step 1), then detecting (Step 2) and reconstructing (Step 3) missing spatial regions, we obtain a sequence of complete, structurally contiguous slices on which the cross-slice 3D-Omics-Flow interpolation (Step 4) can operate without topological distortion.

#### Step 4: Pairwise Generation and Cross-Interval Assembly

Step 4 instantiates the building block Eq. (1) on each adjacent slice pair, fitting the matched-region flow model and the unmatched-region birth–death process to generate intermediate cell states; predictions from adjacent intervals are then stitched together at the observed slice levels to form a single 3D volume. See the later subsection titled “The Two-Slice Building Block: Components and Training” for full details. The in-plane coordinates (*s*_1_, *s*_2_) are returned to their original scale by inverting the standardization used in pre-processing.

##### Generalization to Multiple Slices

For *K* ordered slices *D*_1_, …, *D*_*K*_ stacked along the vertical direction so that 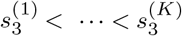, the building block Eq. (1) is applied pairwise on each adjacent interval [*k* − 1, *k*] with its own depth parameter *t* ∈ (0, 1) anchored at the two endpoint slices, and a step count *T*_*k*_ proportional to the estimated number of cell layers in the interval. The interval-specific vertical rescale

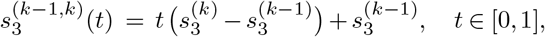

maps the depth parameter to physical *z*-coordinates. Predictions from different intervals meet at the observed slices 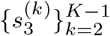; at these vertical levels we retain the observed cells and discard any generated ones to avoid duplication. The post-flow calibration of Step 5 is then applied once globally to the assembled stack.

#### Step 5: Post-Flow Calibration

We calibrate the 3D coordinates produced by the building block Eq. (1) (assembled in Step 4) by aligning them to all observed cell locations (both on the observed omics slices and in an auxiliary form described below) with a 3D cubic B-spline free-form deformation (FFD), yielding calibrated coordinates for downstream visualization and analysis. The auxiliary cell-location information takes one of two forms: (i) additional 2D sections with in-plane locations obtained, e.g., from H&E segmentation, each assigned a vertical coordinate *s*_3_ from its relative position to the observed omics slices; or a 3D volume of cell locations spanning the region between adjacent observed omics slices. The FFD is fitted once globally on the assembled stack.

##### Coordinate Rescale

For the FFD description below, we use **s** = (*s*_1_, *s*_2_, *s*_3_) to denote the 3D coordinates of cells, where the in-plane (*s*_1_, *s*_2_) rescaling and the depth-parameter mapping *t* ↦ *s*_3_ are as described in Step 4 above (when *K* = 2, the depth-parameter mapping reduces to linear interpolation between 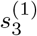 and 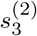.

##### Free-Form Deformation Using Cubic B-Splines

A regular 3D control lattice **P** with *M* control points per axis covers the assembled stack, with grid spacings

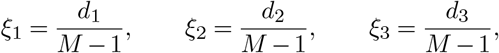

where *d*_*l*_ = *s*_*l*,max_ − *s*_*l*,min_ are the axiswise ranges. The lattice is padded by three control points at each end of each axis to support cubic interpolation at boundaries. Each point **s** is mapped to fractional offsets (*u*_1_, *u*_2_, *u*_3_) ∈ [0, 1)^3^ and lattice indices (*β*_1_, *β*_2_, *β*_3_) via

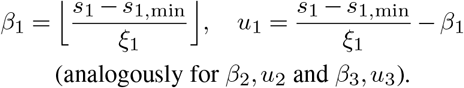

With cubic basis functions

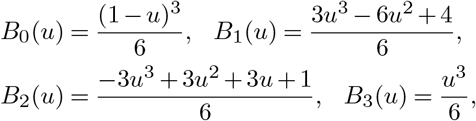

for each source point **s**_*i*_, the deformed position is

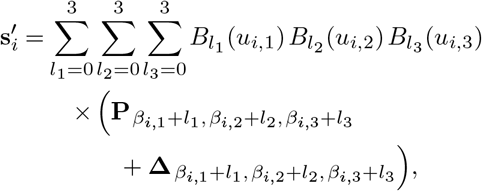

where **Δ** is the array of learnable control-point displacements.

##### Training Objective

We optimize the FFD by minimizing a boundary-weighted Chamfer loss over the control-point displacements **Δ**. Let 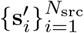 and 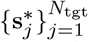 denote the transformed and target coordinates, respectively. The loss is

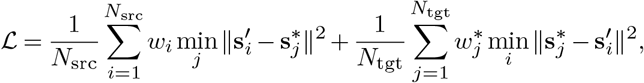

where *w*_*i*_ and 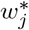 are boundary-proximity weights that assign higher importance to cells near the tissue boundary. Each cell’s weight is computed from its distance *d* to the nearest tissue boundary via an inverse power law on distance, then min-max rescaled to [1, *κ*_*w*_]:

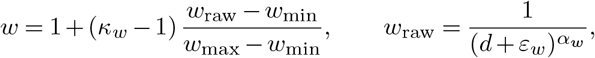

where *κ*_*w*_ *>* 0 controls the maximum up-weighting of boundary cells relative to interior cells (default *κ*_*w*_ = 10), *ε*_*w*_ *>* 0 prevents singularity at the boundary (default *ε*_*w*_ = 1, in the original spatial units), and *α*_*w*_ *>* 0 governs the decay rate (default *α*_*w*_ = 1). The formula is applied separately to source and target weights, with *w*_min_ and *w*_max_ being the minimum and maximum of *w*_raw_ within each side. Both source and target weights are normalized so that ∑_*i*_*w*_*i*_ = *N*_src_ and 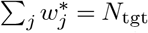, ensuring that the two Chamfer terms remain on comparable scales. For 2D-section input, the segmented cells on each section provide the targets 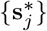 at their measured (*s*_1_, *s*_2_) and the section’s assigned *s*_3_, and in addition to the boundary weighting, we up-weight *w*_*i*_ for source points near each section’s *s*_3_ level; for 3D-volume input, all observed positions are used uniformly as targets. We use Adam with learning rate 0.05 for 100 iterations on a control lattice with *M* = 5 control points per axis.

##### The Two-Slice Building Block: Components and Training

Throughout this subsection, 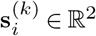 refers to the postalignment, *z*-scored coordinates from Step 1’s Preprocessing; the inverse standardization is applied at the end of the assembly step (Step 4). We use local slice indices *k* ∈ {1, 2} for the two input slices of a single building-block invocation Eq. (1). These correspond to the anchor pair (*k*_−_, *k*_+_) in Step 3 and to an adjacent pair (*k* − 1, *k*) in Step 4. The slice tuples *D*_1_, *D*_2_ and the building-block input/output are as introduced in the Interlude of the previous subsection.

##### Initial Matching and Partitioning

We start by identifying matched cell pairs across two spatial omics slices via an existing matching method (e.g., SLAT (15)). This approach pairs cells across the two slices by jointly leveraging their expression profiles and spatial coordinates. Taking SLAT as a concrete example, in addition to yielding matched cell pairs, it outputs embedding vectors of cells (distinct from the CAST graph embedding **X**^(*k*)^ used as the cell features in the flow model below). Hence, we can compute similarity scores among cells within and between slices based on their embeddings.

Formally, let 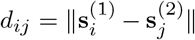 denote the Euclidean distance between the spatial locations of a candidate pair (*i, j*), with *i* on slice 1 and *j* on slice 2, and let cos_*ij*_ denote the corresponding cosine similarity of their SLAT embeddings. The matched pairs are filtered based on two thresholds: *d*_*ij*_ ≤ *τ*_*d*_ and cos_*ij*_ ≥ *τ*_*s*_, where thresholds *τ*_*d*_ and *τ*_*s*_ correspond respectively to the 0.95-quantile of spatial distances and the 0.05-quantile of embedding similarities among all matched pairs outputted by SLAT. Cells that cannot be paired across the two slices, or paired cells that fail filtering, are classified as “unmatched cells”, while cells in matched pairs that pass filtering are classified as “matched cells”. After labeling all cells, we partition the cells of *D*_1_ and *D*_2_ into matched and unmatched subsets: for *k* = 1, 2,

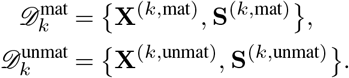

The community and cluster annotations from Step 1 travel with each cell when *D*_*k*_ is partitioned; we suppress them in the tuple notation for brevity. Within the matched subsets, we further re-index cells so that the *i*-th matched cell on slice 1 is paired with the *i*-th matched cell on slice 2.

The interpolation between the matched subsets 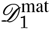 and 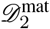 is modeled using the prior-informed flow model, whose representation, training, and prediction we describe next. The interpolation between the unmatched subsets 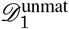 and 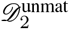 is modeled using the bin-level birth–death process described later in this subsection.

##### Cell State Encoding

The training data for the flow model are cells in the matched regions of the two slices, with 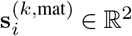 and 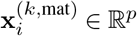 denoting the coordinate and feature vectors for cell *i* in slice *k*. In the rest of the description, we drop “mat” in the superscript for notational simplicity. For cell *i* from slice *k* in the training set, we identify its 20 spatial nearest neighbors from the same slice and define the neighborhood feature as the mean of their features (excluding the cell itself), 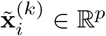. We then define its cell

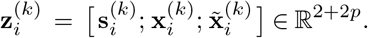

More generally, at depth *t* ∈ [0, 1], we define the state vector analogously,

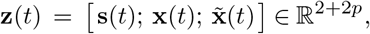

where **s**(*t*) ∈ ℝ^2^, **x**(*t*) ∈ ℝ^*p*^, and 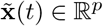 denote, respectively, spatial coordinates, cell features, and neighborhood features at depth *t*.

Conceptually, the trajectory **z**_*i*_(*t*) tracks the transition of matched pair *i* across *t* ∈ [0, 1], with 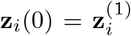 and 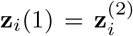. At any intermediate *t* ∈ (0,1), 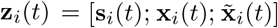 describes a hypothetical cell at depth *t*, with spatial location **s**_*i*_(*t*), feature vector **x**_*i*_(*t*), and neighborhood feature 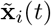. Ideally, all cells along the trajectory should be biologically related, and transitions between consecutive states should respect the structure of the underlying tissue and its biological condition. These principles guide the trajectory modeling that follows.

##### Flow Modeling of Cell State Transition Trajectories

We adapt Rectified Flow (16) and Guided Flow (17) to model smooth state transitions along *t* ∈ [0, 1] between the two end slices. We model these transitions by the following ordinary differential equation (ODE) with boundary conditions:

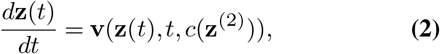

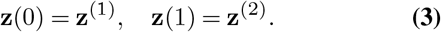

Here, **z**^(1)^ and **z**^(2)^ are the observed states at *t* = 0 and *t* = 1, respectively, defining the boundary conditions of the ODE. In addition, *c*(**z**^(2)^) is the Leiden cluster label of the slice-2 endpoint cell, providing a soft target for where the transition process ends. Throughout, *c* is the cell’s cluster label: a Leiden cluster from Step 1’s Preprocessing by default, or expert cell-type annotation when provided. The cluster centroids used later inherit the same convention. Although the boundary condition **z**(1) = **z**^(2)^ fully specifies the endpoint, conditioning on *c* regularizes the *intermediate* trajectory by encouraging it to follow a velocity field consistent with the target cluster, yielding smoother and more biologically plausible interpolations at *t* ∈ (0, 1). The velocity 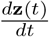 decomposes into three components:

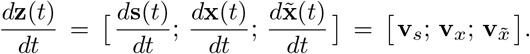

We regard equations Eq. (2)-Eq. (3) as an overarching model for all such trajectories. To learn the velocity function from data, we need to (i) parametrize it in a way that respects tissue structure and cell-type differences, and (ii) identify pairs of endpoints (**z**_*i*_(0), **z**_*i*_(1)) of realized trajectories.

We regularize velocity estimation to respect tissue structure and cell-type differences in two ways. First, by regarding different spatial communities as surrogates for different tissue structures, we require all realized trajectories between matched regions to stay within the same spatial community for all *t* ∈ [0, 1]. Second, we let the velocity function vary across spatial communities and across distinct cell types within each community. Below, we describe how we identify trajectory endpoints and parameterize the velocity function.

##### Within-Spatial-Community Training Pair Curation

We refer to the observed cell states at the endpoints of a trajectory as a training pair. As we require all trajectories between the matched regions to stay within the same spatial community, each pair of endpoints must share a spatial community label. To identify training pairs, we first fix a spatial community *q* ∈ {1,…, *m*_*q*_} and perform linear-sum assignment between the matched cells of community *q* on slices 1 and 2, using the following squared distance. For cell *i* on slice 1 and cell *j* on slice 2, the squared distance is

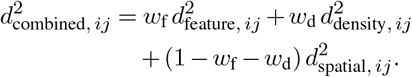

(with default weights *w*_f_ = 0.01 and *w*_d_ = 0.01) where

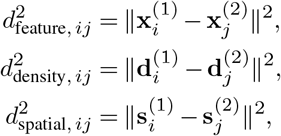

and 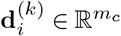 stacks per-cluster cell counts within a fixed spatial radius of 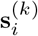, where the radius is set to one tenth of the smaller of the two per-axis coordinate standard deviations on slice *k*. If cell type annotations are not available, we use Leiden cluster labels of the omics features instead.

In practice, the two observed slices may not contain the same number of cells in a given spatial community in the matched regions. To address this, we apply the following augmentation to balance cell counts. Let *n*_*k,q*_ denote the number of cells in the matched region of slice *k* that belong to spatial community *q*. Suppose *n*_1,*q*_ *> n*_2,*q*_. We then sample *n*_1,*q*_ − *n*_2,*q*_ cells with replacement from the *n*_2,*q*_ cells on slice 2. For each randomly selected cell, we apply an independent spatial coordinate perturbation by an *ε*_aug_ sampled from a bivariate 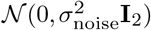 distribution. Here, *σ*_noise_ ≈ *σ*_loc_*/*10 and *σ*_loc_ is the per-axis standard deviation of spatial coordinates of the matched subsets. If *n*_2,*q*_ *> n*_1,*q*_, we apply the foregoing procedure with the roles of the two slices switched. After augmentation, we have *n*_1,*q*_ = *n*_2,*q*_ = *n*_*q*_, and linearsum assignment on the augmented data outputs *n*_*q*_ training pairs.

##### A Spatial-Community-Specific Graph Convolutional Network Parametrization of Velocity

To allow the velocity function to differ across spatial communities, we adopt spatialcommunity-specific parameterization. Within each community, we parametrize a community-specific velocity function **v**_*q*_ via a graph convolutional network (GCN) shared across all trajectories whose endpoints are curated training pairs. Fix a spatial community *q* and obtain the graph structure *G*_*q*_ = (*V*_*q*_, ℰ_*q*_) from all matched cells in community *q* on slice as follows. Each cell forms a node in *G*_*q*_ and there is an edge between two different cells if and only if they are mutual neighbors within a spatial radius (set to the median distance to the 30th nearest neighbor) and mutual *κ*_*f*_-nearest neighbors in feature space, with *κ*_*f*_ = 30 by default (distinct from Step 1’s *κ*, which sets the within-slice spatial graph). At each node, the input is (**z**, *t, c*), where *c* ∈ {1,…, *m*_*c*_} is the Leiden cluster label (*m*_*c*_ = total number of Leiden clusters), encoded as a one-hot vector in 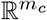 for input to the GCN. Note that the endpoint cluster label *c* stays unchanged along each trajectory. The input has four components: spatial coordinates **s** concatenated with *t*, cell features **x**, neighborhood features 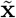, and cluster label *c*. Each is first mapped through a separate fully-connected encoder to a 256dimensional hidden representation. The four representations are concatenated and passed through a fully-connected layer with output dimension 256, then two graph convolution layers (256 →256 → 2*p* + 2), all with hyperbolic tangent activation. We write the foregoing parametrization as

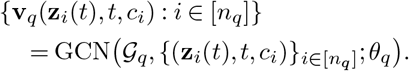

Here, *n*_*q*_ is the number of training pairs curated for spatial community *q*, and *θ*_*q*_ collects the weight matrices used at different layers of the GCN; *θ*_*q*_ alone constitutes the spatialcommunity-specific trainable velocity parameters.

##### Prior-Informed Training Objective

Within a spatial community *q*, we train the GCN-based flow model by minimizing a composite loss function ℒ_flow_:

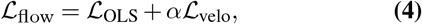

where *α* is a hyperparameter weighting the two terms (default *α* = 1.0).

The ordinary least squares (OLS) loss is:

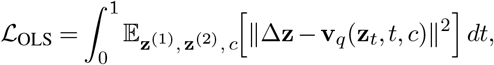

where Δ**z** = **z**^(2)^ − **z**^(1)^, and **z**_*t*_ = (1 − *t*)**z**^(1)^ + *t***z**^(2)^ denotes the linear interpolation between the two boundary states (distinct from the ODE trajectory **z**(*t*)). Here, the expectation is over curated training pairs in community *q*. This OLS loss corresponds to the standard rectified flow objective, which encourages straight-line interpolation paths between boundary states, so that the learned ODE produces approximately linear trajectories between slices.

The velocity regularization loss ℒ_velo_ encourages feature velocities to align with cell-type transition directions. Specifically, we perform a new Leiden clustering of cell features on all cells spatial community *q* on the two slices. Let the centroids of the resulting Leiden clusters be 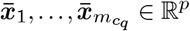. We define the following set of canonical cell feature transition directions:

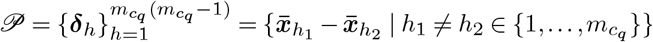

This set provides a reference for plausible velocity directions, and we define ℒ_velo_ as:

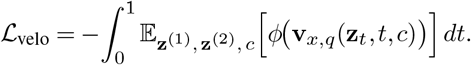

where **v**_*x,q*_ ∈ ℝ^*p*^ denotes the feature-velocity component of **v**_*q*_, and

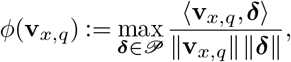

where for any vectors **a** and **b**, ⟨**a, b**⟩ denotes their inner product.

##### Other Training Details

During training, we apply condition dropout by randomly replacing *c* with a null symbol ∅ in each training pair with probability *p*_drop_, as a regularizer that prevents over-reliance on the cluster label:

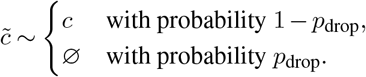

where ∅ is an all-zero vector in 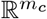. By default, *p*_drop_ = 0.1 for all communities. This dropout is applied only during training; at inference, we use *c* directly. We use the Adam optimizer to minimize ℒ_flow_ in Eq. (4), with learning rate 5 × 10^−3^ and 1000 training iterations per community; each iteration performs one Adam step on the gradient computed over all curated training pairs in the community.

##### Generative Sampling Algorithm

After training, we have learned a velocity function **v**_*q*_ for each spatial community *q*. Within each spatial community *q*, given the cell state of the endpoint of each training pair on slice 1 and the cluster label of the paired endpoint on slice 2, we can now generate the unobserved trajectories for all curated training pairs by numerically integrating the velocity **v**_*q*_.

We adopt the Euler method to generate each trajectory iteratively:

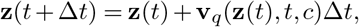

where Δ*t* = 1*/T*, and *T* is the estimated number of cell layers between the two observed end slices, assuming each cell layer is approximately 10*µ*m thick. We denote each generated trajectory by {**z**_fwd_(*t*) : *t* ∈ (0, 1)}. As *t* goes from 0 to 1, this trajectory moves from slice 1 to slice 2.

To place the two observed slices on equal footing, we swap the roles of **z**^(1)^ and **z**^(2)^ in model building and training and obtain another generated trajectory {**z**_bwd_(*t*) : *t* ∈ (0, 1)} that evolves in the backward direction; as *t* goes from 0 to 1, the backward trajectory moves from slice 2 to slice 1. The symmetric combination of both trajectories yields the final generated trajectory between each curated training pair:

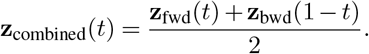

To respect the original cell counts within each spatial community *q*, we apply the following thinning procedure to the combined trajectories. Let *n*_1,*q*_ and *n*_2,*q*_ denote the number of matched cells in community *q* on slices 1 and 2, respectively. At each generated time step *t*, the target count is *n*_*q*_(*t*) = (1 − *t*) *n*_1,*q*_ + *tn*_2,*q*_ (rounded to the nearest integer). If *n*_1,*q*_ ≥ *n*_2,*q*_ (shrinking community), we group the combined trajectories by their endpoint on slice 2; among trajectories sharing the same endpoint, each is independently terminated with probability 1 − *n*_*q*_(*t*)*/n*_*q*_(*t* − Δ*t*) at step *t*, so that the expected surviving count matches *n*_*q*_(*t*). If *n*_1,*q*_ *< n*_2,*q*_ (expanding community), we reverse direction: group by endpoints on slice 1 and apply the analogous thinning from *t* = 1 backward.

Repeating the foregoing generation across all *m*_*q*_ spatial communities in the matched regions of the two observed slices, we have now predicted the unseen volume sandwiched between the matched regions.

##### Bin-Level Birth–Death Process for Unmatched Regions

For the unmatched subsets 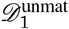 and 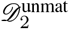 on the two observed slices, we model the spatially varying cell population dynamics via a stochastic birth–death process with spatially smoothed, per-cluster rates inferred from observed cell count changes between the two slices.

##### Setup and Bin-Level Counts

We partition the spatial domain into bins of side length 50 *µ*m, indexed by *r*. Let *c* index the Leiden cluster labels obtained from Step 1’s Preprocessing. For slice *k* ∈ {1, 2}, define the bin–cluster count

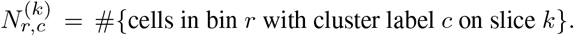

Let 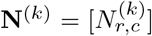 denote the resulting matrix for slice *k* where each row corresponds to a bin and each column to a Leiden cluster label.

##### Birth–Death Rate Estimation

For each (*r, c*) we estimate per-capita birth and death rates 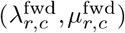 from **N**^(1)^ to **N**^(2)^ under a no-immigration constraint (i.e., neither rate can create cells in bins where 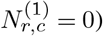. With a small *ε*_bd_ *>* 0,

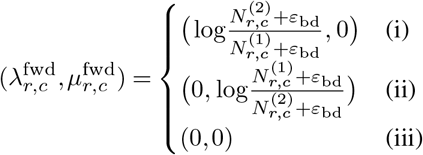

with default *ε*_bd_ = 10^−5^, where the three cases are

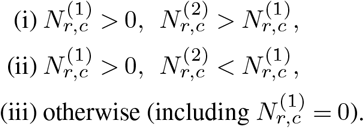

Backward per-capita birth and death rates 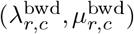 are obtained by exchanging the roles of slice 1 and slice 2:

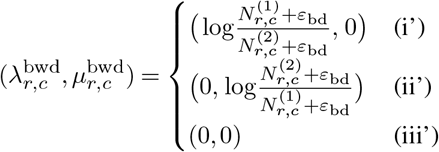

where the three cases are

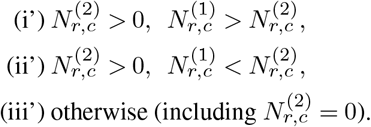

##### Spatial Smoothing of Rates

To stabilize rate estimates across neighboring spatial bins, we smooth both forward and backward per-capita rates over a bin-level *κ*_bin_-nearestneighbor graph. Let 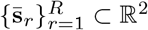 denote the bin centers (computed as the mean of cell coordinates within each bin). Construct a binary adjacency matrix **W** ∈ {0, 1}^*R×R*^ with **W**_*rr*′_ = 1 if *r*′ is among the *κ*_bin_ nearest neighbors of *r* (excluding *r*) and **W**_*rr*′_ = 0 otherwise (default *κ*_bin_ = 4). Define the row-normalized adjacency matrix 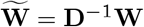 with **D**_*rr*_ = ∑_*r*_*′* **W**_*rr*′_. For a smoothing strength *η* ∈ [0, 1] (default *η* = 0.2) and each cluster label *c*, update the forward and backward birth–death rates by

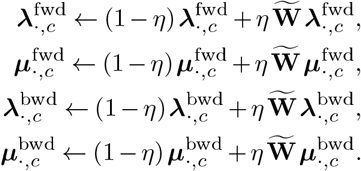

This update is applied once.

For notational simplicity, unless otherwise stated, we use (*λ, µ*) to denote the smoothed birth and death rates hereafter.

##### Interpolation of Cell Counts within Bins

Given the smoothed birth–death rates (*λ*_*r,c*_, *µ*_*r,c*_), we first sample *N*_*r,c*_(*t*), the bin-level cell count for cluster *c* in bin *r* at an intermediate time *t*. We discretize the time interval [0, 1] into *T* equal steps of length Δ*t* = 1*/T*, where *T* is estimated from the number of cell layers between the two observed end slices. At each time step, we sample birth–death events as follows:

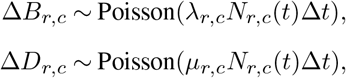

The population count is then updated at each step according to:

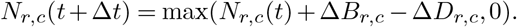

A single Poisson realization is drawn per direction at each step. Symmetry is restored by averaging the forward and backward trajectories defined below. For *t* ∈ [0, 1], let **N**_fwd_(*t*) be the evolution of **N**^(1)^ to time *t* under forward rates as defined above. By switching the roles of the two slices and using backward rates, we define **N**_bwd_(*t*) analogously. We form the final cell count interpolation as

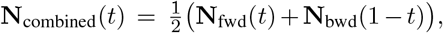

with overrides in the following special cases: if 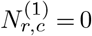 and 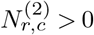, set *N*_combined, *r,c*_(*t*) = *N*_bwd, *r,c*_(1 − *t*); if 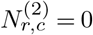 and 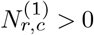, set *N*_combined, *r,c*_(*t*) = *N*_fwd, *r,c*_(*t*).

##### Sampling Cell Coordinates and Features

For each *t*, given **N**_combined_(*t*) we sample cells from a reference pool 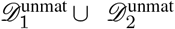 stratified by (*r, c*) with replacement. Specifically, we first partition the reference cells by bin *r* and cluster label *c*. Then, for each stratum (*r, c*), we draw exactly *N*_*r,c*_(*t*) cell indices uniformly at random from that stratum, allowing repeats when *N*_*r,c*_(*t*) exceeds the number of available reference cells in (*r, c*) (if a stratum is empty, no cells are drawn for it). For sampled cells we: (i) take feature vectors from the embedding **X**^(*k*)^ used elsewhere in the pipeline and add small Gaussian noise (default standard deviation 10^−3^); (ii) perturb spatial coordinates by isotropic Gaussian noise of per-axis standard deviation *σ*_loc_ *t/*10, where *σ*_loc_ is the smaller of the two per-axis coordinate standard deviations within the bin and *t* ∈ [0, 1] is the depth parameter (so the noise grows linearly with depth). Combining the matched-region trajectories **z**_combined_(*t*) with the unmatched-region samples at each *t* yields the virtual slice *D*^∗^(*t*), comprising its feature matrix **X**^∗^(*t*) and coordinates **S**^∗^(*t*), which together realize the building-block mapping in Eq. (1).

### Benchmark Design and Evaluation

#### Train–Validation Split and Leakage Control

For each benchmarking experiment, the input slice stack is partitioned into training slices (used for matching, flow-model fitting, and bin-level rate estimation) and held-out validation slices (used only for evaluation). Per-dataset splits:

- *Fig. 1 (CyCIF Human Colorectal Cancer, natural damage)*. Three training slices: slice25 (*z* = 125 *µ*m, intact anchor), slice39 (*z* = 195 *µ*m, carrying natural tissue damage and a simulated E-cadherin dropout), and slice54 (*z* = 270 *µ*m, intact anchor). Two heldout validation slices: slice34 (*z* = 170 *µ*m) and slice44 (*z* = 220 *µ*m).
- *Fig. 2 (CyCIF Human Colorectal Cancer, combined damage)*. Eleven training slices spanning ~460 *µ*m along *z*: slice14, slice20, slice25, slice34, slice44, slice54, slice64, slice74, slice84, slice97, slice106 at *z* = 70, 100, 125, 170, 220, 270, 320, 370, 420, 485, 530 *µ*m, respectively. Three held-out validation slices: slice59 (*z* = 295 *µ*m), slice78 (*z* = 390 *µ*m), slice102 (*z* = 510 *µ*m).
- *OpenST Human Lymph Node (Supplementary Fig. S5– S6)*. Five training slices: slice 2 (*z* = 58 *µ*m, intact), slice 9 (*z* = 261 *µ*m, damaged), slice 17 (*z* = 493 *µ*m, damaged), slice 26 (*z* = 754 *µ*m, intact), and slice 34 (*z* = 986 *µ*m, intact); damage on slice 9 and slice 17 is spatial only (no feature dropout). Four held-out validation slices: slice 5 (*z* = 145 *µ*m), slice 11 (*z* = 319 *µ*m), slice 23 (*z* = 667 *µ*m), and slice 28 (*z* = 812 *µ*m).
- *INSIHGT Mouse Hypothalamus (Supplementary Fig. S2–S4)*. Ten evenly spaced training slices (slice1, slice9, slice17, slice25, slice33, slice41, slice49, slice57, slice65, slice73) spanning *z* = 12–732 *µ*m, and nine midpoint validation slices (slice5, slice13, slice21, slice29, slice37, slice45, slice53, slice61, slice69). No damage is applied; the dataset is used to assess clean-input fidelity.

Validation cells are excluded from every stage of model fitting (alignment, matching, training). Competitor parity is enforced by construction: SpatialZ and the linear interpolation baseline are fitted on exactly the same training slices with identical preprocessing, and only then are predictions extracted at each validation *z*.

#### Synthetic Damage Simulation

Synthetic Damages are constructed by overlaying two perturbations onto training slices: marker dropout (setting one or more feature columns to NaN) and tissue loss (removing all cells inside a 2D mask). For Fig. 1 the tissue loss is not simulated — the validation target slice carries pre-existing natural damage, and only the marker dropout is synthetic.

a. *Marker dropout*. For Fig. 1, E-cadherin is set to NaN on slice39. For Fig. 2, Ki67 and *α*-SMA are jointly set to NaN on slice20, slice34, slice54, and slice74; E-cadherin is set to NaN on slice25, slice44, slice64, and slice84. OpenST Human Lymph Node and INSIHGT Mouse Hypothalamus receive no feature dropout. In every experiment dropout is applied to **Y**^(*k*)^ before spatial alignment, biomarker restoration, and embedding.
b. *Tissue-loss masks*. On each damaged slice, cells whose (*x, y*) fall inside a fixed coordinate region are removed from **Y**^(*k*)^ and **S**^(*k*),raw^. Tissue loss masks in Fig. 2 are: *Group A* = slice20 with *x* ∈ [5000, 10000], *y* ∈ [10000, 20000]; *Group B* = slice34 and slice44 with *x* spanning the full slice extent and *y* ∈ [16000, 20000]; *Group C* = slice74 and slice84 with *x* ∈ [15000, 25000], *y* ∈ [15000, 20000]. Eleven training slices fall into three categories: *intact* (slice14, slice97, slice106), *feature-dropout only* (slice25, slice54, slice64), and *combined damage* carrying both spatial removal and feature dropout (slice20, slice34, slice44, slice74, slice84). For OpenST Human Lymph Node, *x* ∈ [2000, 10000], *y* ∈ [4000, 6000] is removed on slices 9 and 17. Damage simulation involves no random sampling, the damaged inputs are deterministic.

#### Benchmarking Methods: SpatialZ and Linear Interpolation Baseline

We compare 3D-Omics-Flow with two competitors: the published method SpatialZ (5) and an in-house linear interpolation baseline. All methods are fit using the same training slices and evaluated at the same held-out validation *z* depths. Neither baseline includes tissue-loss detection or restoration.

a. *SpatialZ baseline*. Because SpatialZ does not accept missing features, marker dropout is first imputed using the corresponding feature mean computed across the training slices. We run SpatialZ’s `Generate_multiple_slices` routine using the training slices ordered by their *z* coordinates. SpatialZ generates virtual slices between each consecutive pair of anchor slices; the number of intermediate slices per pair is set to the estimated number of cell layers between the two anchors, computed using the same procedure as in 3D-Omics-Flow (the cell-layer count *T* defined in “Interpolation of Cell Counts within Bins”). Each virtual slice is assigned a linearly interpolated *z* coordinate between its two parent anchors. For each validation depth, we take the SpatialZgenerated slice whose interpolated *z* coordinate matches the target depth as the prediction.
b. *Linear interpolation baseline*. For each validation depth *z*_target_, we identify the two flanking training slices *k* and *k*+1, where *z*_*k*_ *< z*_target_ *< z*_*k*+1_, and compute

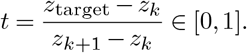

Cells across the two anchor slices are matched by spatial proximity, and each matched pair is linearly interpolated in both spatial coordinates and biomarker expression. Specifically, each matched pair (*i, j*) yields a virtual cell

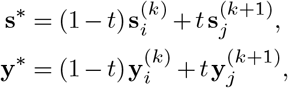

where **s** denotes spatial location and **y** denotes the raw biomarker feature vector. As with SpatialZ, missing feature values caused by simulated marker dropout are mean-imputed from the training data, and no tissue-loss restoration is performed.

#### Reconstruction Quality Metrics (Fig. 1, CyCIF Human Colorectal Cancer small stack)

Quality of the validation-slice reconstruction is reported with three metrics, all computed on the in-plane (*s*_1_, *s*_2_) at the validation *z* depth and compared against the ground-truth slice.

1. *Cell-type-normalized L*_1_ *similarity (Fig. 1e)*. At grid bin size *b* ∈ {10, 20,…, 100} *µ*m, the validation slice is partitioned into square bins indexed by *r*. For each bin and cell type *c* ∈ {1,…, *C*}, let 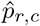 and 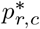 be the cell-type composition vectors in the reconstruction and ground truth, normalized so that 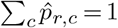 and 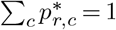 within each nonempty bin (per-bin normalization across cell types). Let ℛ (*b*) denote the union of bins that are non-empty in either the reconstruction or the ground truth at bin size *b* (bins empty in both are dropped). The reported similarity is

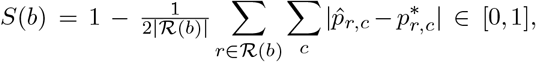

which is higher when reconstruction and ground-truth compositions agree more closely. Fig. 1e plots *S*(*b*) averaged across the two validation slices.
2. *Feature cosine similarity (Fig. 1f)*. Features are the *raw* biomarker columns shared between the reconstruction and the ground truth. Each marker is robust-scaled to [0, 1] across cells, after which the per-bin mean feature vector 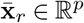 is the column-wise mean of the scaled markers over cells in bin *r*. For each bin *r* ∈ ℛ (*b*) we compute the cosine similarity cos 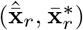, and report the bin-level mean

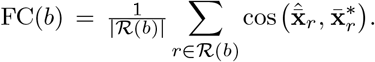
3. *Per-marker MS-SSIM (Fig. 1g)*. Each marker channel of the validation slice is rasterized to an *n* × *n* image whose pixel grid spans the joint (*x, y*) bounding box of the reconstruction and the ground truth, with per-pixel value equal to the mean expression of cells falling in that pixel; the default reported resolution is *n* = 1024. Each rasterized marker is robust-scaled to [0, 1] (0.5%–99.5% percentile clip), lightly pre-smoothed with `scipy.ndimage.gaussian_filter` (*σ* = 0.2 pixels), and min–max normalized to [0, 1]. The score between reconstruction and ground truth is then computed with `skimage.metrics.structural_similarity`. Per-marker scores are averaged across the two validation slices; Fig. 1g reports the per-marker difference (3D-Omics-Flow) − (SpatialZ) in purple and (3D-Omics-Flow) − (Linear) in gray.

#### Tumor–Immune Invasive Margin Zone Analysis (Fig. 2, CyCIF Human Colorectal Cancer large stack)

To quantitatively interrogate the volumetric tumor microenvironment recovered by 3D-Omics-Flow on the CyCIF Human Colorectal Cancer dataset, we label the reconstructed volume into five biologically meaningful zones and profile cellular composition, marker expression, and tissue morphometry within each zone.

#### Zone Definition via Signed Distance Field

Tumor cells (Tumor/Epi., Ki67+, PDL1+) are voxelized at 25 *µ*m to form a binary tumor mask, which is dilated by a 3 × 3 × 3 structuring element to close pixel-scale gaps. The signed distance field (SDF), computed by Euclidean distance transform on the dilated mask, satisfies SDF(**s**) *<* 0 inside the tumor and 0 outside, in microns. Each cell is mapped to its enclosing voxel and assigned a zone label by its SDF and immune status:

- *Core tumor*: SDF *<* −*d*_in_ (deep inside the tumor mass).
- *Margin tumor*: −*d*_in_ ≤ SDF *<* 0 (tumor cells at the invasive front).
- *Immune at margin*: immune-lineage cells with −*d*_in_ ≤ SDF *< d*_out_.
- *Margin stroma*: non-immune cells with 0 ≤ SDF *< d*_out_.
- *Distal*: all remaining cells.

Default thresholds are *d*_in_ = 50 *µ*m and *d*_out_ = 100 *µ*m.

#### 3D Spatial Architecture Metrics

Panels k–o of Fig. 2 quantify the recovered 3D tumor microenvironment using the invasive-margin zones and signed distance field defined in “Tumor–Immune Invasive Margin Zone Analysis” above.

1. *Mean zone expression (Fig. 2k)*. Each marker is *z*-scored across all cells of the 3D-Omics-Flow 3D reconstruction; for each (zone, marker) pair, the mean *z*-score over cells in that zone is reported. Four invasive-margin zones (*core tumor, margin tumor, immune at margin, margin stroma*) are shown.
2. *Nearest tumor* → *immune distance (Fig. 2l)*. A *margin-tumor* cell is a cell with margin_zone = margin_tumor, and a *margin-immune* cell is a cell with margin_zone = immune_at_margin. For each margin-tumor cell we record the 3D Euclidean distance (in *µ*m) to its nearest margin-immune cell, and pool the distances across all margin-tumor cells to form the reported distribution.
3. *Voxel-level marker correlation (Fig. 2m)*. The 3D-Omics-Flow reconstruction is voxelized on the same 25 *µ*m 3D grid used for zone definition (spacing 25 *µ*m in *x, y, z*). For each occupied voxel and each marker, the per-voxel value is the mean expression over the cells whose (*x, y, z*) falls in that voxel; voxels with zero cells are dropped (not zero-filled). For every pair of markers we compute the Pearson correlation across the common set of occupied voxels.
4. *Tumor–immune contact z-score (Fig. 2n)*. The analysis is restricted to *interface cells* (margin_zone ∈ margin_tumor, immune_at_margin). For each tumor subtype *T* and immune subtype *I*, the observed contact value is the mean number of interface immune cells of subtype *I* within 60 *µ*m of an interface tumor cell of subtype *T*,

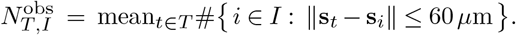

A null distribution is generated by shuffling immune-subtype labels among interface immune cells while holding all other quantities fixed; *n*_perm_ = 200 permutations are drawn, and the reported score is

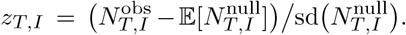
5. *Marker-vs-distance gradient (Fig. 2o)*. To recover continuous variation along the tumor-normal direction (which discrete zone means collapse), we use the signed distance field as a signed micron coordinate assigning each cell a unique position relative to the tumor boundary (negative inside, positive outside). The SDF axis is partitioned into uniform 10 *µ*m bins spanning [ −100, +200] *µ*m, covering the dense tumor core, the invasive margin, and the proximal stroma; we plot the bin-averaged expression of every marker against this axis, with SDF = 0 marking the tumor boundary.

#### 2D vs. 3D Margin-vs-Core Analysis (Supplementary Fig. S7)

To assess whether 3D reconstruction improves the recovery of biological signal, we first label zones of two reference, 2D sparse training stack before reconstruction and 3D-Omics-Flow reconstruction dense bulk. In both settings, zone labels were then transferred to held-out validation slices (slice59, slice78, and slice102) by nearest-neighbor lookup. Specifically, for each validation cell, we identified the closest labeled reference cell in Euclidean space using only spatial coordinates and assigned the corresponding zone label. This procedure uses only reference coordinates, reference labels, and validation coordinates; validation biomarker measurements are not used and therefore cannot leak into the zone assignment. For downstream analysis, we compared *core_tumor* and *margin_tumor* cells in the validation slices. For each marker in the panel {Ki67, *α*-SMA, Ecadherin, PD1, PDL1, CD3, CD20, CD68, Keratin, CDX2, PCNA, NaKATPase, LaminABC}, we applied Welch’s two-sided *t*-test using `scipy.stats.ttest_ind` followed by the Holm–Bonferroni adjustment for multiple testing using `statsmodels.stats.multitest.multipletests`. We also computed the log_2_ fold change of the group means, 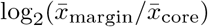. The resulting test significance and effect size define the volcano plot in Supplementary Fig. S7b, with the dashed reference line indicating *p*_adjusted_ = 0.05.

### 3D Cell–Cell Communication and Marker Domain Analysis (Supplementary Fig. S4, INSIHGT Mouse Hypothalamus)

On the INSIHGT Mouse Hypothalamus dataset, we assess two aspects of biological fidelity that per-slice 2D metrics cannot evaluate: preservation of spatially co-localized ligand–receptor signaling and preservation of spatially coherent multi-marker domains. Both analyses operate on the full 3D-Omics-Flow-reconstructed 3D volume and compare against the ground-truth 3D dataset.

#### Cell–Cell Communication Score

To probe whether ligand– receptor co-localization is preserved in 3D, we curate twelve biologically motivated marker pairs spanning GABAergic, cholinergic, catecholaminergic, neuropeptidergic, glial, and neuronal-structural axes (e.g., GABA–GABRG2, TH– DBH, SST–NPY, GFAP–AIF1, TUBB3–MAP2, Orexin A– HTR3A); scoring each pair in both directions yields 24 directed ligand–receptor tests. For each cell *i* we build a 3D radius-neighbor graph at *r* = 60 *µ*m on cell coordinates, row-normalize its adjacency matrix, and compute the per-cell interaction score

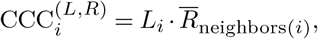

the product of the cell’s own ligand expression and the mean receptor expression among its 3D neighbors — a juxtacrine/paracrine co-expression signal that depends on true 3D adjacency rather than within-slice 2D proximity. Per-cell scores are computed independently for ground-truth and 3D-Omics-Flow reconstructions and compared as overlaid histograms.

#### Marker Domain Analysis

To test whether spatially coherent 3D domains of marker co-expression are preserved, the 25 markers are grouped into six biologically curated functional sets: GABAergic (GABA, GABRG2, PVALB, CALB1, CALB2, SST, VIP, NPY), cholinergic (CHAT), catecholaminergic (TH, DBH), glial (GFAP, CNP1, AIF1), neuronal-structural (TUBB3, MAP2, RBFOX3, RBFOX2), and neuropeptide (Orexin A, Galanin, GLP1, HTR3A).

Within each dataset every marker is *z*-score standardized; for each cell and group we then take the per-cell domain score to be the mean *z*-score across that group, a composite that captures coordinated activity of a functional axis while suppressing single-marker noise. Per-cell domain score distributions are compared group by group between ground-truth and 3D-Omics-Flow reconstructions, reading out whether the reconstruction preserves prevalence and dynamic range for each functional program in 3D.

## Code and data availability

3D-Omics-Flow and related tutorials are freely available to the public at GitHub: https://github.com/MaBLD/3DOmicsFlow/. Data and Code to regenerate the main and supplementary figures are also deposited to GitHub.

## ACKNOWLEDGEMENTS

We thank Dr. HeiMing Lai for providing us with 3D volumetric images before they became available. The work is supported by the Parker Institute for Cancer Immunotherapy (G.P.N.), and the Rachford and Carlota A. Harris Endowed Professorship (G.P.N.), the NIH HuBMAP consortium grant U54HG012723 (Y.T., M.S., and G.P.N), PROMISE PROJECT-2025-27 SCI-W grant (Y.T., G.P.N.), the NIH HTAN grant U01CA294514 (Z.M.), and the NSF grant DMS-2245575 (Z.Z. and Z.M.). This article reflects the authors’ views and should not be construed as representing the views or policies of the NIH or other institutions that provided funding.

## AUTHOR CONTRIBUTIONS

Conceptualization: Z.Z., Y.T., G.P.N., Z.M.

Methodology: Z.Z., Z.M.

Data Analysis and Benchmark: Z.Z., Y.T., Z.M.

Data Resources and Valuable Inputs: Y.T., M.S., G.P.N.

Original Draft: Z.Z., Y.T., Z.M.

Supervision: M.S., G.P.N., Z.M.

