## Supplementary Figures for "Fault-tolerant 3D reconstruction from 2D spatial proteomics sections"

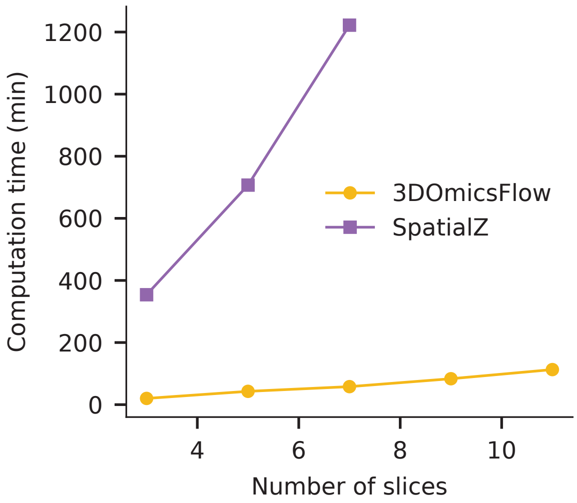


**Supplementary Figure S1: Runtime comparison between 3D-Omics-Flow and SpatialZ on the CRC proteomics dataset.** End-to-end wall-clock time as a function of the number of input slices (3, 5, 7, 9, 11), with two adjacent consecutive slices spaced $\sim$50 $\mu$m apart in $z$ and $\sim$20,000 cells per slice. Both 3D-Omics-Flow and SpatialZ grow near-linearly with depth, while SpatialZ scales much more steeply.

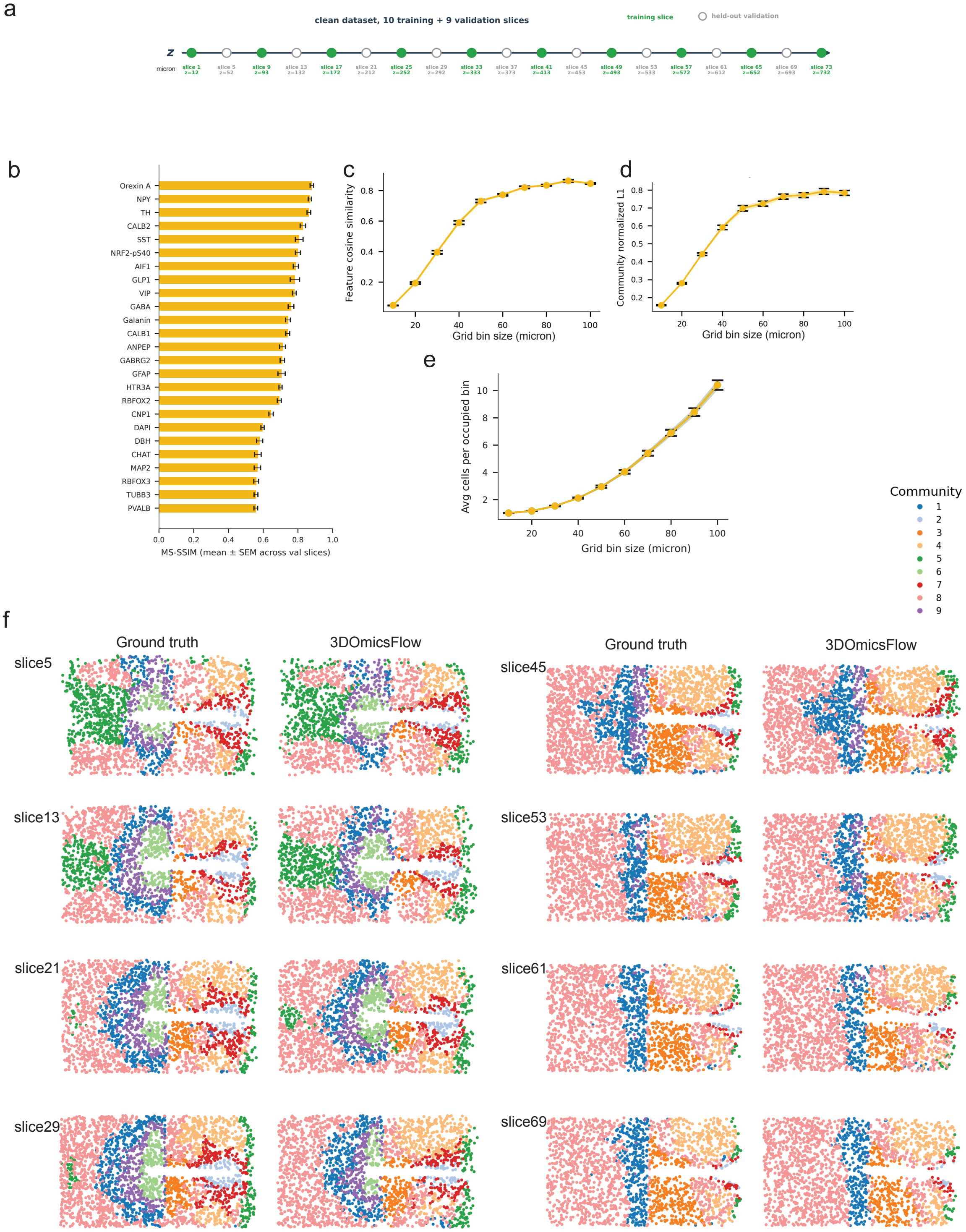


**Supplementary Figure S2: Validation of 3D-Omics-Flow on a clean, densely sampled real 3D INSIHGT Mouse Hypothalamus dataset (8). a,** Experimental design: 37 serial slices spanning $z = 12\text{–}732\,\mu$m of an undamaged spatial-transcriptomics tissue; 10 evenly spaced slices (green) are used for training and the 9 midpoint slices between consecutive training pairs (open circles) are held out for validation. **b,** Per-feature MS-SSIM between 3D-Omics-Flow predictions and ground truth on the 9 validation slices (mean $\pm$ s.e.m. across slices; 25 features). **c–e,** Spot-level metrics on validation slices as a function of grid bin size (10–100 µm): feature cosine similarity (**c**), community-normalized L1 (**d**; higher is better) and the average number of cells per occupied bin (**e**). Mean $\pm$ s.e.m. over the 9 validation slices. **f,** Spatial scatter plots of held-out validation slices (9 communities); ground truth and 3D-Omics-Flow are shown side-by-side, demonstrating recovery of community geometry across the full $z$-range.

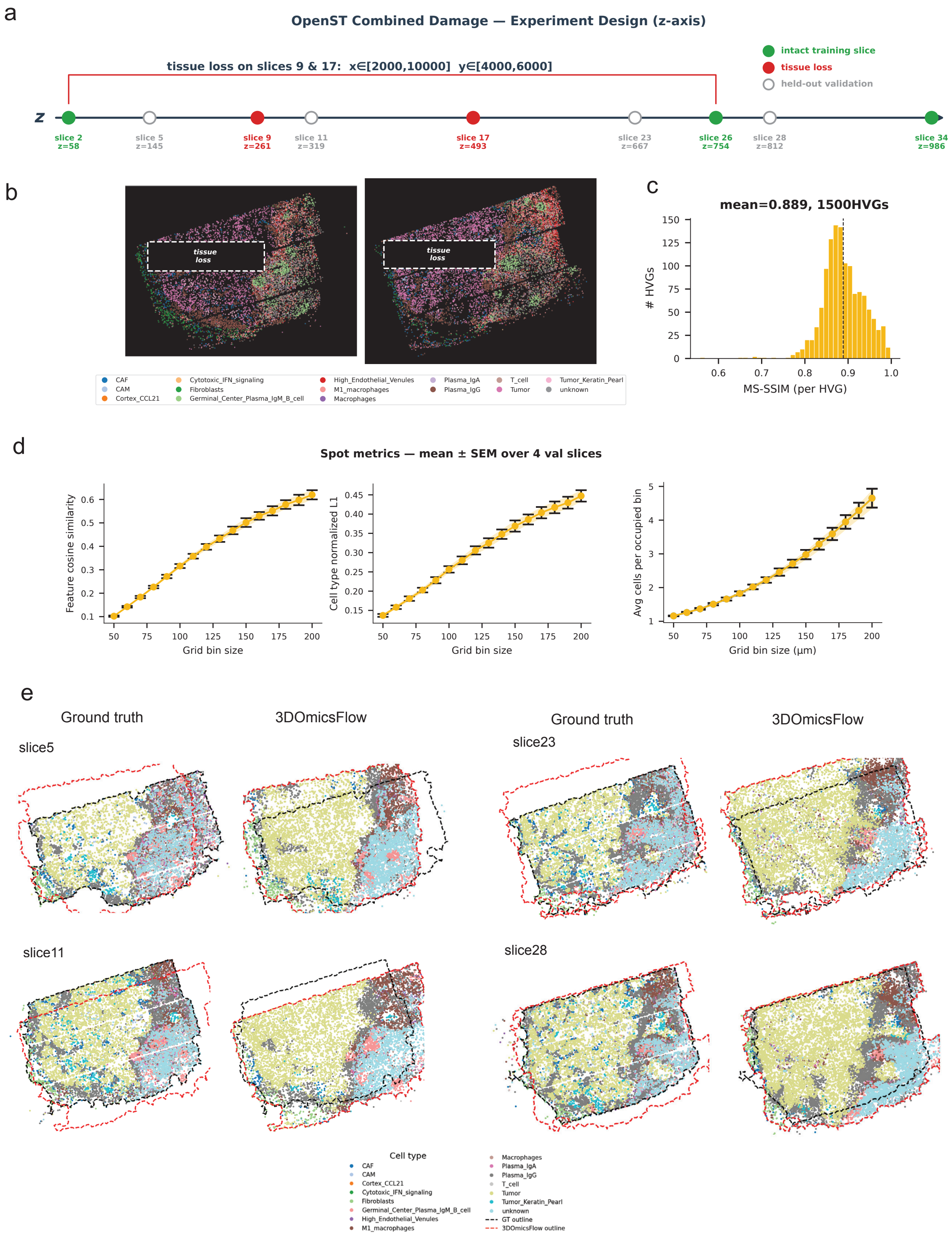


**Supplementary Figure S3: 3D-Omics-Flow generalizes to a transcriptomics dataset (OpenST) with simulated spatial damage. a,** OpenST experimental design: 5 training slices (2, 9, 17, 26, 34) span the $z$-axis; slices 9 and 17 carry rectangular tissue loss ($x \in [2000, 10000]$, $y \in [4000, 6000]$). Open circles, validation slices interleaved between training slices. **b,** Spatial scatter of the two damaged training slices (9 and 17) colored by cell type; dashed white box indicates the simulated missing region. **c,** Distribution of per-HVG MS-SSIM between 3D-Omics-Flow predictions and ground truth on the held-out slices (1,500 highly variable genes; mean MS-SSIM $= 0.889$). **d,** Spot-level metrics on validation slices vs grid bin size (50–200 µm): feature cosine similarity, cell-type normalized L1 and average cells per occupied bin. Mean $\pm$ s.e.m. across 4 validation slices (5, 11, 23, 28; one per inter-training gap). **e,** Spatial plots of cell types across all 4 held-out validation slices, ground truth (left) vs. 3D-Omics-Flow prediction (right). For each section, the red boundary outlines the 2D section boundary of the 3D volume predicted by 3D-Omics-Flow at the $z$-coordinate of the held-out validation slice, the boundary of which is outlined in black. The difference between the boundaries is due to large $z$-axis gaps across adjacent training slices (**a**). We focus on overlapping regions of ground truth and the prediction by 3D-Omics-Flow across validation slices.

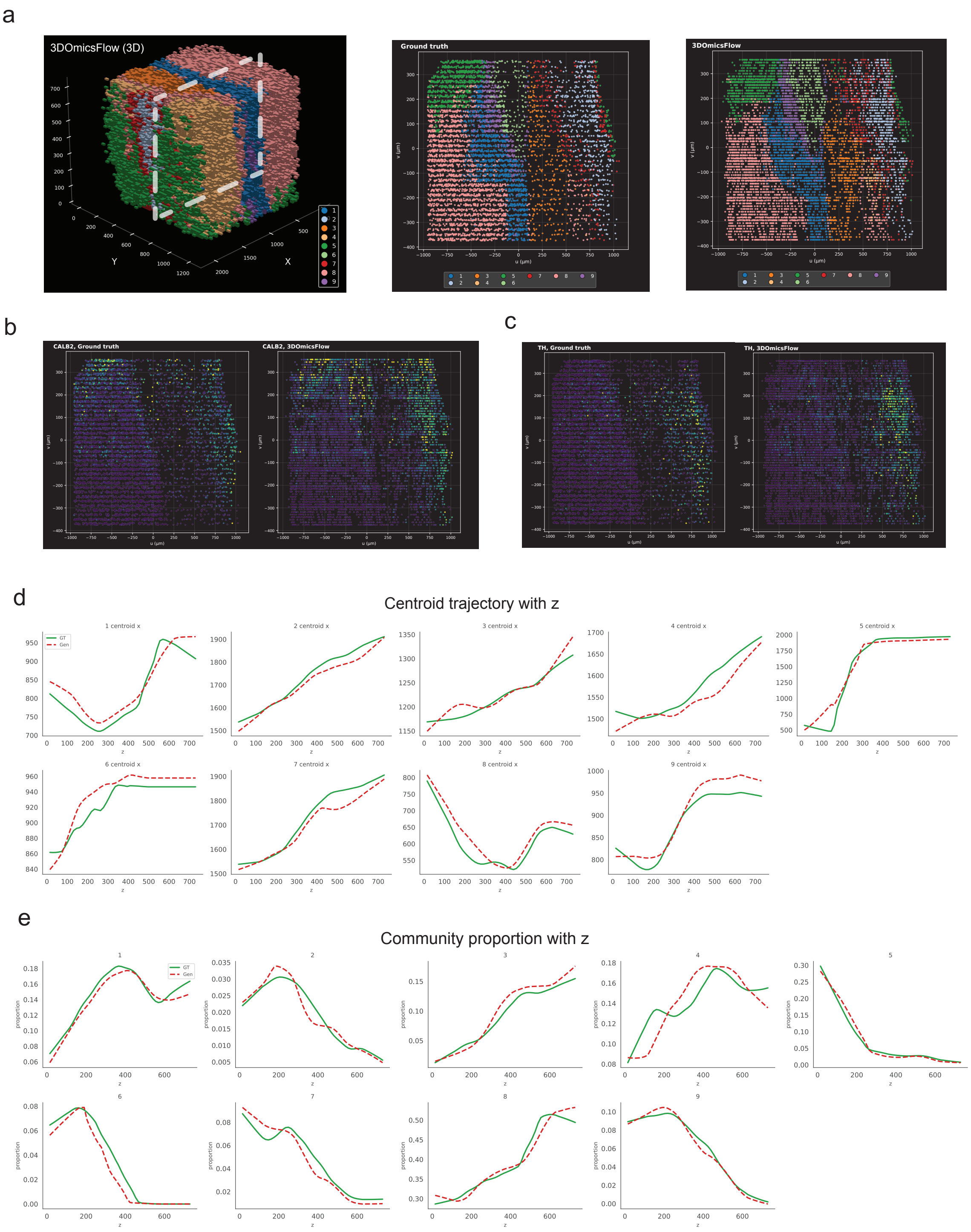


**Supplementary Figure S4: 3D structural fidelity of the 3D-Omics-Flow reconstruction on the INSIHGT Mouse Hypothalamus dataset (8). a,** Left, full 3D-Omics-Flow 3D reconstruction; the dashed white plane marks the oblique slab used in the right two panels. Middle and right, in-slab 2D scatter for ground truth vs 3D-Omics-Flow. **b,c,** Single-feature views of the same slab (ground truth vs 3D-Omics-Flow), colored by scaled expression. **d,** Per-community spatial centroid trajectories along $z$: each panel plots the $x-$ centroid of one community as a function of $z$ for ground truth (red) and 3D-Omics-Flow (green). **e,** Per-community proportion as a function of $z$ (fraction of cells in each $z$-bin assigned to that community), ground truth (red) vs 3D-Omics-Flow (green).

a

| Domain | Markers | Biology |
|---|---|---|
| GABAergic | GABA, GABRG2, PVALB, CALB1, CALB2, SST, VIP, NPY | Inhibitory interneuron system |
| Cholinergic | CHAT | Acetylcholine-producing neurons |
| Catecholaminergic | TH, DBH | Dopamine/norepinephrine neurons |
| Glial | GFAP, CNP1, AIF1 | Astrocytes, oligodendrocytes, microglia |
| Neuronal_structural | TUBB3, MAP2, RBFOX3, RBFOX2 | Pan-neuronal structural markers |
| Neuropeptide | Orexin A, Galanin, GLP1, HTR3A | Neuromodulatory peptides |

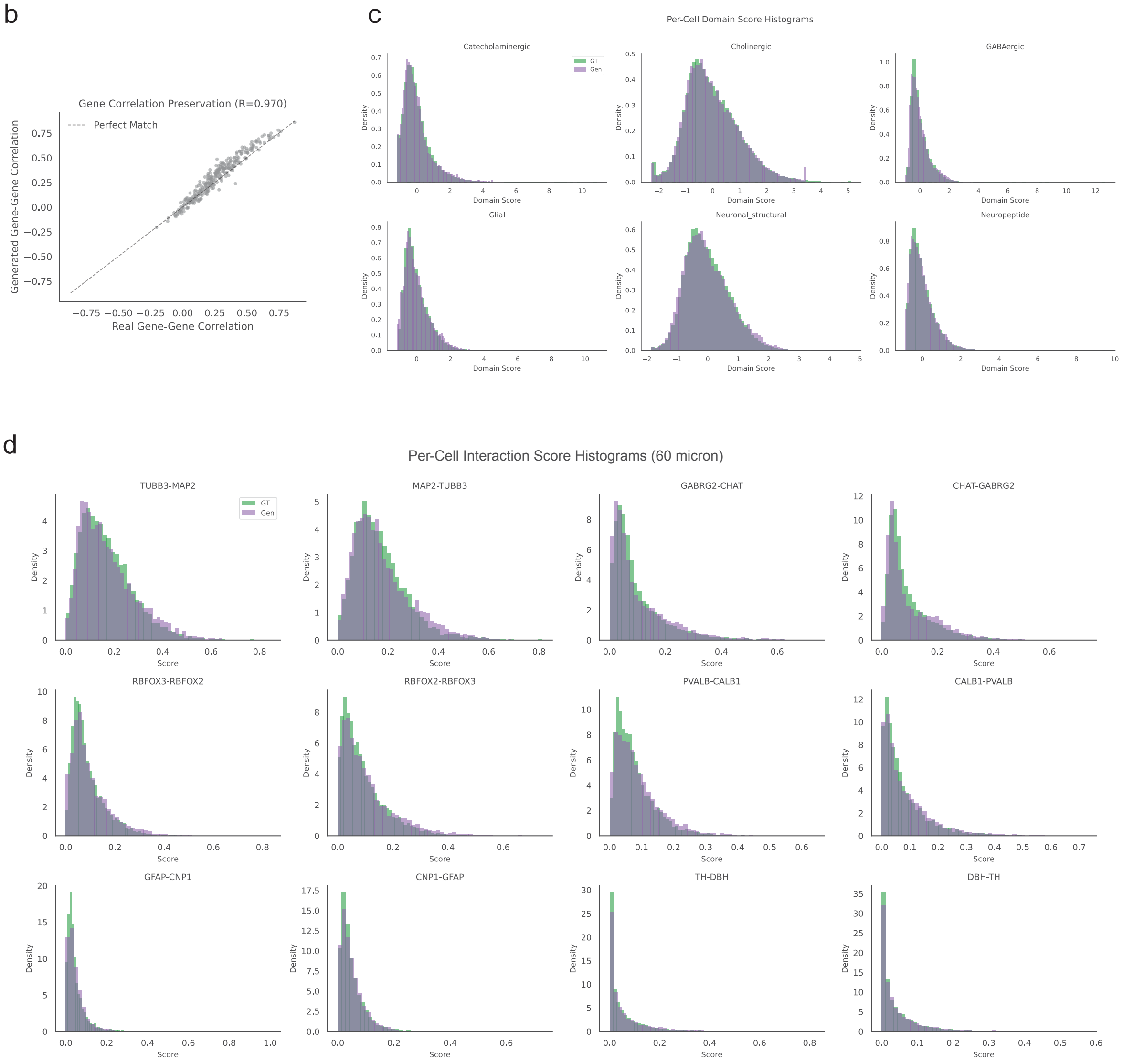


**Supplementary Figure S5: 3D molecular structure preservation: cell–cell communication and marker-domain analysis (INSIHGT Mouse Hypothalamus dataset (8)). a,** Six biologically curated marker domains used for downstream domain analyses: GABAergic, Cholinergic, Catecholaminergic, Glial, Neuronal_structural and Neuropeptide, with their constituent markers and biological role. **b,** Marker–marker correlation preservation. Pairwise Pearson correlations between all 25 markers are computed independently in ground truth and 3D-Omics-Flow; each dot is one marker pair. Dashed line, $y = x$; overall $R = 0.970$. **c,** Per-cell domain score distributions for each of the six marker groups. Domain score = mean of $z$-scored marker expression within the group; histograms compare ground truth (green) and 3D-Omics-Flow (purple). **d,** Per-cell cell–cell interaction (CCC) score distributions for 12 directed ligand–receptor / marker pairs at a 3D neighborhood radius of 60 µm. Score per cell $i$: $L_i \times \overline{R}_{\mathrm{neighbors}(i)}$, where neighbor expression is averaged on a row-normalized radius graph in 3D, with each dataset independently rescaled to $[0, 1]$ per feature (0.5–99.5 percentile). Ground truth (green) vs 3D-Omics-Flow (purple).

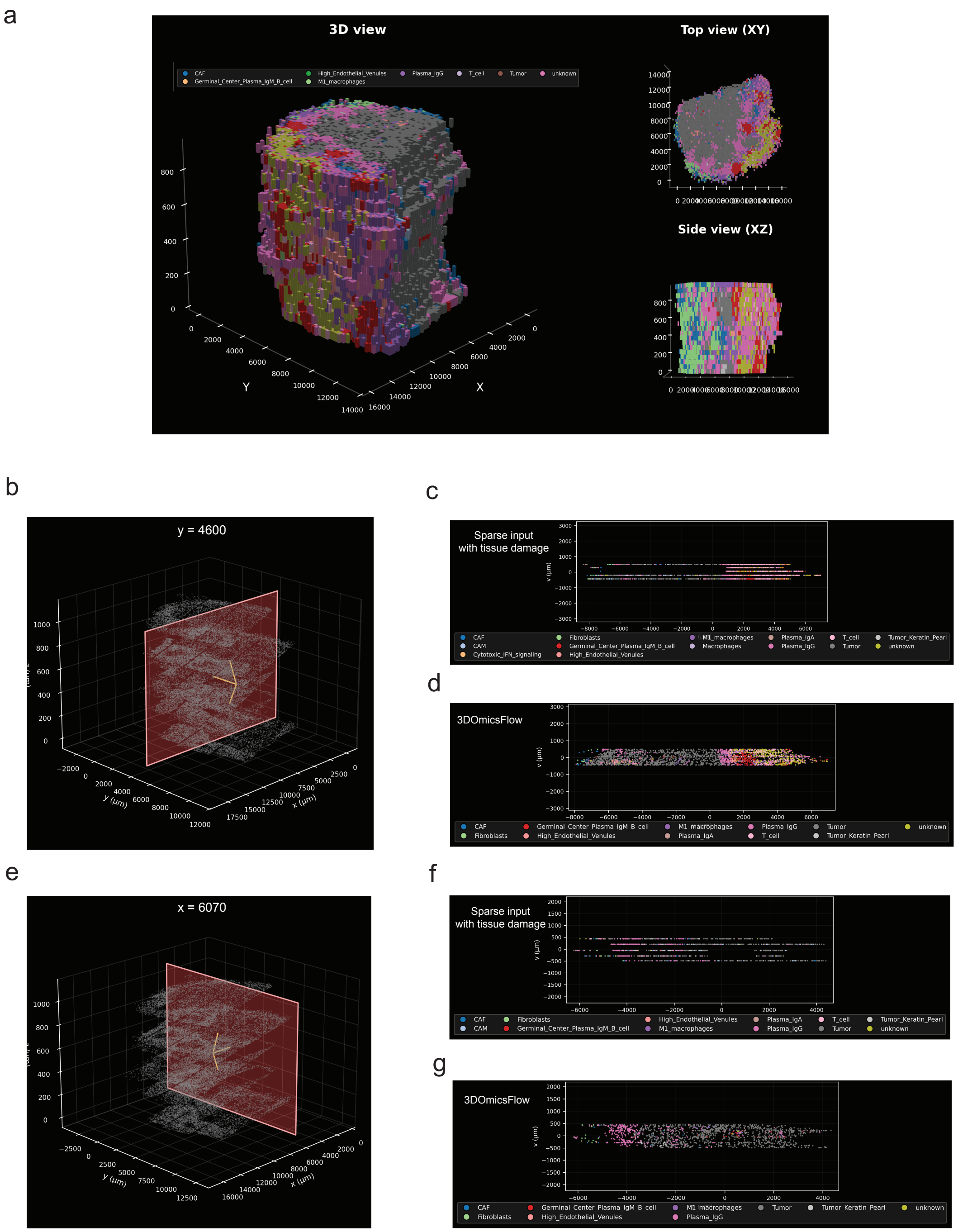


**Supplementary Figure S6: 3D structure of the OpenST reconstruction and recovery of damaged tissue regions. a,** 3D-Omics-Flow reconstruction of the OpenST volume rendered at 25 µm voxel resolution, colored by cell type. Left, oblique 3D view; top right, XY top-down view; bottom right, XZ side view. **b,** XZ context view of the full reconstructed volume (light gray) with a sagittal cutting plane at $y \approx 4000$ (red) intersecting the damage region. **c,d,** 2D in-slab scatter projected onto the plane in **b**, colored by cell type: damaged sparse input (**c**) vs the 3D-Omics-Flow reconstruction (**d**); cells lost in the input box are restored after reconstruction. **e,** As in **b** but with an orthogonal cutting plane at $x \approx 6000$, also passing through the damage centroid. **f,g,** 2D in-slab scatters for the cut in **e**: damaged input (**f**) vs 3D-Omics-Flow (**g**).

a

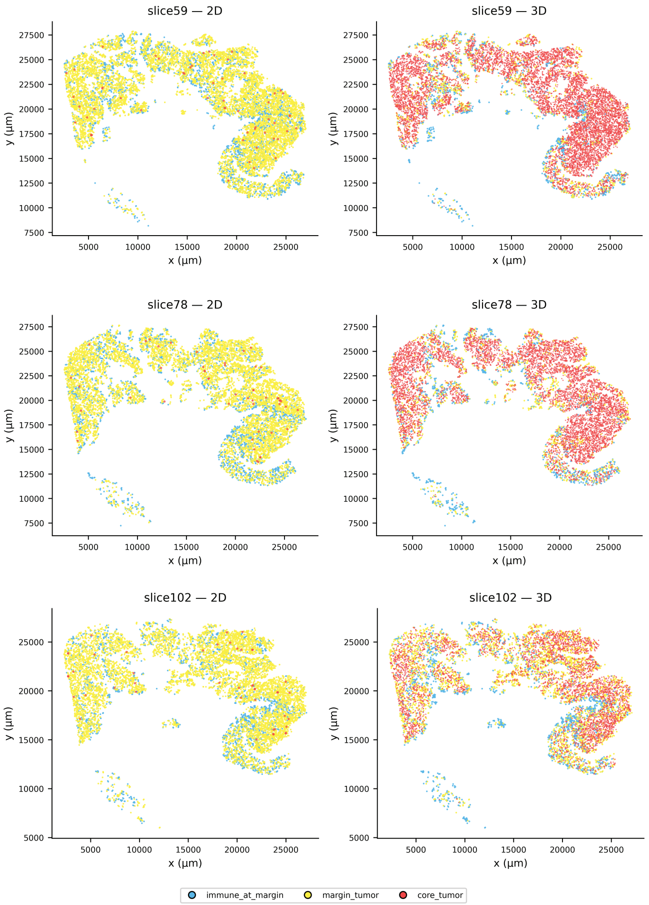


b

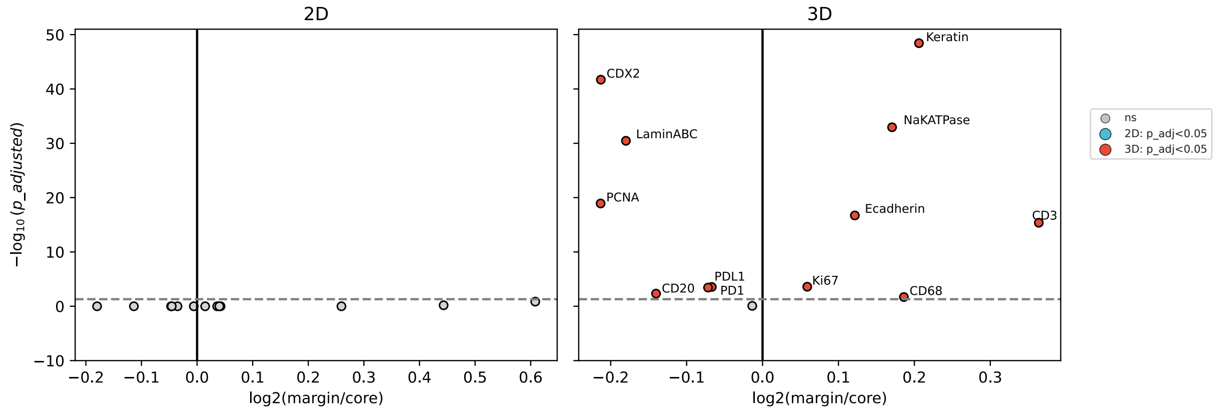


**Supplementary Figure S7: 3D reconstruction recovers tumor–immune zonation and statistical structure that sparse 2D sections cannot resolve.** Validation slices (slice59, slice78, slice102) are held out from training, so their cells receive zone labels by nearest-neighbor transfer from already-zoned cells in two independent reference pools. *2D-on-sparse:* reference consists of the sparse 2d training sections after zonation labeling; *3D-on-dense:* reference consists of the 3D-Omics-Flow-reconstructed dense volume. Validation-cell biomarker values used in test statistic and $P$-value calculations in panel **b** are not used at any step of zone assignment. **a,** Per-slice spatial plots of tumor cells on the three held-out validation slices (rows: slice59, slice78, slice102), colored by projected zone (core_tumor, red; margin_tumor, yellow; immune_at_margin, cyan). Left column: 2D-on-sparse projection; right column: 3D-on-dense projection. The 3D projection resolves a coherent core/margin geometry that the sparse 2D projection cannot. **b,** Markers volcano plot of margin-tumor vs core-tumor cells on the held-out validation slices (slices 59, 78, and 102; cf. **Fig. 2a**). $x$-axis: $\log_2$(mean margin / mean core); $y$-axis: $-\log_{10}$ adjusted p-value (Holm–Bonferroni adjusted p-value from two-sample t-test). Left: zones from the 2D-on-sparse projection; right: zones from the 3D-on-dense projection. The 3D projection uncovers significant margin/core differences for canonical tumor-lineage (Keratin, Ecadherin, NaKATPase), proliferation (Ki67), immune (CD3, CD20, CD68), and immune-checkpoint markers (PD1, PDL1) that the 2D projection fails to detect.
